# CpG islands are topological sinks for transcription-induced DNA supercoiling

**DOI:** 10.64898/2026.08.24.746546

**Authors:** Catherine Naughton, Andrea Bonato, Michael Chiang, Samuel Corless, Jon Stocks, Ewan Forsyth, Graeme R Grimes, David Halliday, Alessandro Bentivoglio, Chris A Brackley, Davide Marenduzzo, Nick Gilbert

## Abstract

Transcription-induced DNA supercoiling is relaxed by topoisomerases, yet negative supercoiling accumulates at active gene promoters. How promoter sequences and chromatin architecture organise and regulate this torsional stress remains poorly understood. Using Twist-seq, a genome-wide assay for DNA supercoiling, we show that highly transcribed CpG island (CGI) promoters accumulate elevated levels of negative supercoiling compared to highly transcribed non-CGI promoters. Coarse-grained molecular dynamics simulations predict that the capacity of CGI promoters to accommodate elevated negative supercoiling arises from the intrinsic physical properties of CGI DNA. CGI sequence and local chromatin architecture favour localised melting at AT-rich pockets and DNA writhe within promoter-proximal nucleosome-free regions, providing an energetically favourable mechanism to dissipate torsional stress. Consistently, CGI promoters show a depletion in Top1 and increased ssDNA in vivo. We propose that CGIs function as sequence-encoded topological sinks that buffer transcription-induced torsional stress, with increased somatic mutation rates at highly expressed CGI promoters suggesting a potential genome-stability cost.

## Introduction

Chromatin structure is highly dynamic and tightly regulated ensuring that essential genomic regions are accessible to cellular machinery when required. Once accessible to transcription, replication, or DNA repair machinery, the DNA duplex undergoes cycles of unwinding and rewinding, rendering it susceptible to topological stress and structural transitions collectively referred to as DNA supercoiling^1–5^. Under normal physiological conditions, the conformational structure of DNA is B-DNA where the two strands are wound around each other in a right-handed helix with ∼10.5 bp per turn of the helix. DNA supercoiling refers to the shift from this relaxed DNA state to one that is either overwound (positive supercoiling) or underwound (negative supercoiling). This phenomenon can be described using two related but distinct structural parameters: twist and writhe^2^. Twist refers to changes in the number of base pairs per helical turn, while writhe describes the spatial coiling of the DNA double helix in space and together these define the overall topology of supercoiled DNA.

In eukaryotes most DNA supercoiling is introduced by transcription as RNA polymerase II (Pol II) generates overwinding of the double helix (positive DNA supercoils) ahead of the transcription complex and leaves underwound DNA (negative supercoils) in its wake^1,6^. This DNA supercoiling is modulated by topoisomerase enzymes that introduce transient breaks in the DNA to relieve topological strain and preserve genomic structural integrity^7–11^. Whilst we have previously shown that negative DNA supercoils, generated by Pol II, accumulate at the transcription start sites (TSSs) of active genes^12^ our understanding of the mechanisms that define the topology and dynamics of supercoiled DNA at gene promoters, and how these are influenced by promoter sequence and architecture, remain poorly understood. To address this, we adapted our psoralen-based protocol for next-generation sequencing (NGS), developing Twist-seq for high-resolution, quantitative profiling of DNA supercoiling genome-wide

In humans, Pol II initiates transcription of protein-coding and non-coding genes from a diverse set of promoters, which can be broadly divided into two categories^13^. CpG island (CGI) promoters, comprising ∼70% of all promoters, are enriched in CpG dinucleotides, typically exhibit broad transcription start sites, and are often associated with ubiquitously expressed housekeeping genes ^14^. In contrast, non–CpG island (non-CGI) promoters make up the remaining ∼30%, frequently include TATA-box promoters with sharp, well-defined start sites and are generally linked to tissue-specific, tightly regulated genes. CGI and non-CGI promoters also differ in their chromatin organisation, with the distinctive sequence composition of CGI promoters contributing to reduced nucleosome occupancy^15^. Intriguingly, previous *in vitro* studies have demonstrated that the structural response of DNA to torsional stress is influenced by its underlying sequence^16^. Together, these differences in promoter sequence and chromatin architecture, coupled with the sequence-dependent response of DNA to torsional stress, led us to ask whether CGI and non-CGI promoters differ in how they accommodate transcription-induced DNA supercoiling.

Here, we present the first high-resolution map of DNA supercoiling at human gene promoters. We reveal a striking and previously unrecognised difference in DNA topology between CGI and non-CGI promoters, with CGI promoters exhibiting substantially elevated negative supercoiling. By integrating genome-wide supercoiling profiling with coarse-grained molecular dynamics simulations and ssDNA-seq analysis, we show that this behaviour is encoded by the intrinsic physical properties of CGI DNA sequence and promoter chromatin architecture. The GC-rich, nucleosome-depleted CGI context concentrates torsional stress onto embedded AT-rich pockets, driving localised DNA melting and facilitating the formation of promoter-proximal topological sinks. We propose that this mechanism buffers transcription-induced torsional stress at highly active genes, potentially contributing to the evolutionary conservation of CGI promoter architecture.

## Results

### Twist-seq quantitatively profiles DNA supercoiling at gene promoters

Previously we developed a psoralen-based method to investigate the dynamics of DNA supercoiling in living cells^12^. This technique employs cell-permeable biotinylated 4,5,8-trimethylpsoralen (bTMP), which preferentially intercalates into underwound DNA, providing a readout of DNA supercoiling across a continuum of topological states, from negative to positive supercoiling^17^. Twist-seq is a NGS-compatible adaptation of this approach that enables high-resolution, genome-wide profiling of DNA supercoiling (Fig. 1a). Briefly, RPE1 cells were incubated with bTMP (500 µg/ml), followed by UV crosslinking of bTMP-bound DNA. DNA was fragmented by sonication, purified, and bTMP-bound DNA was isolated and processed into sequencing-ready libraries. Sequencing read depth per RPE1 sample was a minimum of 200 million reads (PE150), with four combined replicates yielding >1.5 billion reads, corresponding to an average genomic sampling density of approximately one fragment per 1.8 bp. Given the global incorporation of bTMP, this substantial read depth was necessary to achieve the dense sampling required for quantitative profiling of DNA supercoiling at transcription start sites (TSSs) within human gene promoters. Topology-independent effects of bTMP were corrected for by normalising to bTMP-bound genomic DNA.

**Figure 1:**
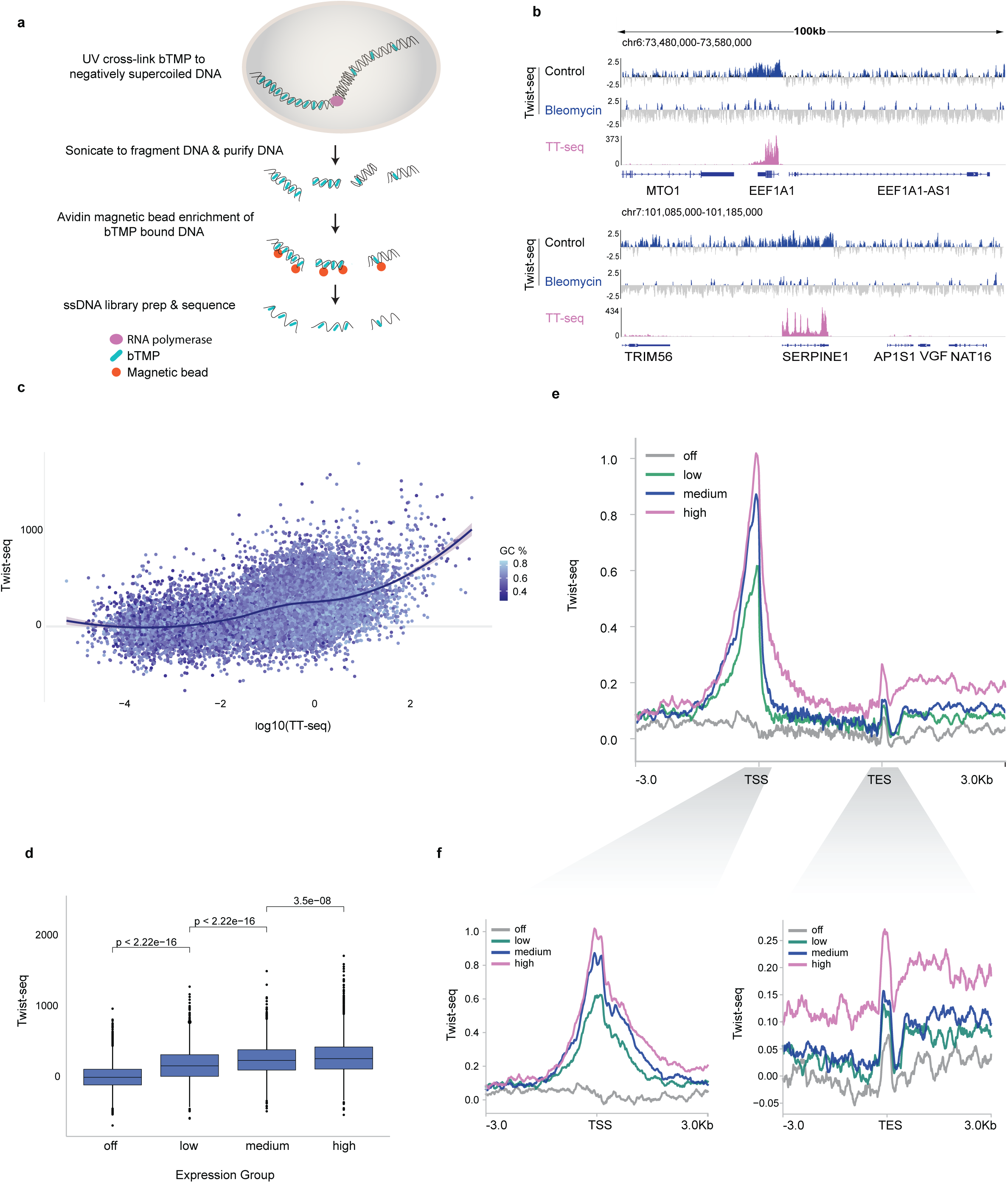
Negative DNA supercoils accumulate at transcriptionally active gene promoters. **a,** Twist-seq protocol: bTMP treatment of cells for 20 min, 360-nm UV cross-linking of bTMP into DNA helix, DNA purification, streptavidin-coated magnetic bead enrichment for biotin-TMP bound DNA, elution of bTMP-DNA (ssDNA), ssDNA specific next gen library preparation and Illumina sequencing to map DNA supercoiling at high resolution genome wide. **b,** Twist-seq mapped genomic distribution of negative DNA supercoiling +/− Bleomycin treatment, around transcriptionally active genes EEF1A1 and SERPINE1. Control (4 replicates) (see Fig. S1a) and Bleomycin (2 replicates). Twist-seq data is log2(Twist-seq control/Twist-seq genomic) in 1 bp bins. TT-seq mapped nascent gene expression is shown as RPKM. **c,** Correlation between the distribution of DNA supercoiling (Twist-seq +/−500 bp at TSS) and transcription (TT-seq, average RPKM/gene) at gene promoters. **d,** DNA supercoiling (Twist-seq RPKM +/− 500 bp at TSS) distribution across expression groups (TT-seq data). *P* values were calculated using nonparametric Wilcoxon two-sided tests. **e,** DNA supercoiling levels (log2(Twist-seq control/Twist-seq genomic)) averaged across all genes, grouped by expression level, along with a zoomed-in view at the TSS and TES with a 3-kb extension on both sides.

Our analysis revealed a significant enrichment of bTMP binding at transcriptionally active genes (e.g. EEF1A1 and SERPINE1) (Fig. 1b and Fig. S1a). The introduction of DNA strand breaks via bleomycin treatment abrogated this signal, highlighting the substantial transcription-induced torsional stress generated at these loci by Pol II activity. Intriguingly there was a threshold level of gene expression beyond which negative DNA supercoils accumulate at gene promoters, which may be indicative of the maximum topoisomerase activity at TSSs (Fig. 1c). To investigate this relationship further genes were split into four quartiles based on nascent RNA expression and found to have a strong correlation between gene expression and promoter bTMP enrichment, consistent with increased negative DNA supercoiling at the promoters of highly transcribed genes (Fig. 1d-f). In agreement with previous observations^18^, these genes were associated with high levels of Pol II binding. A peak of bTMP enrichment was also observed at transcription end sites (TESs), coincident with Pol II pausing prior to release (Fig. S1b and Fig. 1e-f).

### CGI promoters accumulate and buffer elevated negative supercoiling

To investigate how promoter sequence and architecture influence DNA supercoiling we categorised genes into the two main Pol II promoter subtypes: CGI and non-CGI (Fig. 2a). In agreement with prior studies^14^, CGI genes had significantly higher gene expression than non-CGI genes (Fig. S2a). Twist-seq analysis revealed a striking difference in DNA supercoiling at CGI versus non-CGI TSSs indicating that the promoters of constitutively expressed genes experience elevated levels of torsional stress (Fig. 2b). Notably, this DNA supercoiling was GC sequence independent (Fig. S2b). Pronounced bTMP enrichment at CGI promoters remained evident when genes were further classified by expression level (Fig. S2c), consistent with elevated negative DNA supercoiling at CGI promoters across expression levels. (Fig. 2c– e and Fig. S2d).

**Figure 2:**
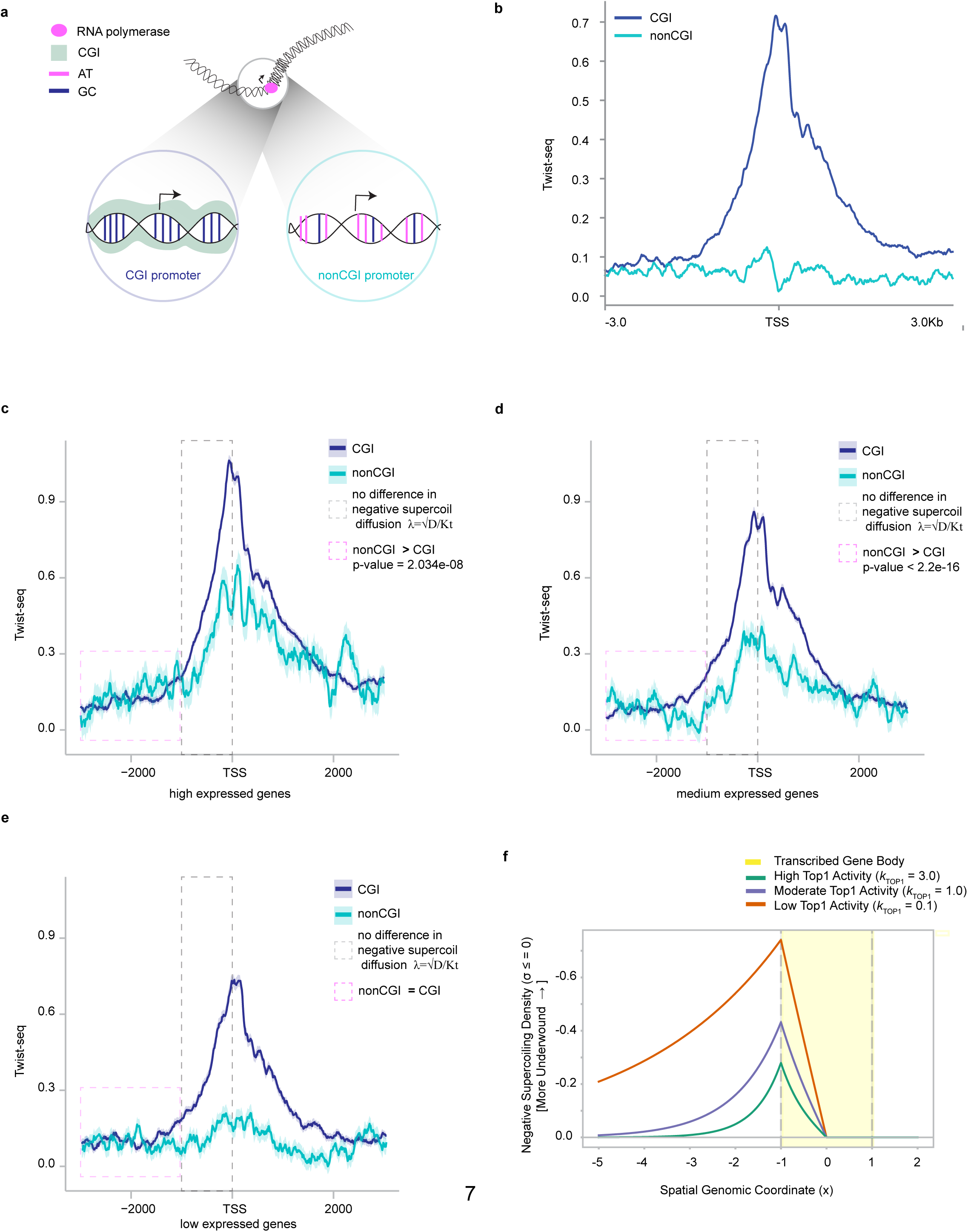
CGI promoters accumulate and buffer elevated negative supercoiling. **a,** Model to explore how negative DNA supercoils behave at CGI versus non-CGI promoters. **b,** DNA supercoiling levels (Twist-seq, 1 bp bins) ±3 kb around TSSs for CGI and non-CGI genes. **c,** DNA supercoiling levels (Twist-seq, 1 bp bins) ±3 kb around TSSs, averaged across all genes within the high expression group for CGI and non-CGI promoters. Statistical significance was evaluated using a two-sided Welch’s t-test. **d,** DNA supercoiling levels (Twist-seq, 1 bp bins) ±3 kb around TSSs, averaged across all genes within the medium expression group for CGI and non-CGI promoters. Statistical significance was evaluated using a two-sided Welch’s t-test. **e,** DNA supercoiling levels (Twist-seq, 1 bp bins) ±3 kb around TSSs, averaged across all genes within the low expression group for CGI and non-CGI promoters. **f,** One-dimensional reaction–diffusion model of transcription-generated DNA supercoiling around the TSS. Steady-state negative supercoiling density (σ ≤ 0) is shown as a function of genomic position for low (kTOP1 = 0.1), moderate (kTOP1 = 1.0) and high (kTOP1 = 3.0) TOP1 relaxation activity. Increasing TOP1 activity reduces both the magnitude and spatial extent of negative supercoiling surrounding the transcribed gene body.

Importantly, amongst highly expressed genes, mean expression was comparable between CGI and non-CGI genes (Fig. S2c,e), yet CGI promoters exhibited substantially greater bTMP enrichment, with area-under-the-curve (AUC) analysis revealing >20% excess bTMP signal (Fig. 2c), consistent with a more negatively supercoiled promoter state. Notably, despite comparable transcriptional output, CGI promoters exhibited higher levels of engaged Pol II than non-CGI promoters (Fig. S2f,g)^19^.

Surprisingly, despite the elevated negative supercoiling at CGI promoters, the TSS-proximal negatively supercoiled domain was similar in size (∼3 kbp) between CGI and non-CGI genes (Fig. 2c). Moreover, upstream of the TSS, levels of negative supercoiling were equivalent or slightly greater at non-CGI promoters (Fig. 2c–e). A reaction–diffusion model of transcription-generated supercoiling showed that changes in supercoil relaxation alter both the magnitude and characteristic spatial extent of the supercoiling domain (Fig. 2f). The increased accumulation of negative supercoiling within a similarly sized domain at CGI promoters therefore suggests that additional promoter-specific properties influence the local accommodation of torsional stress. Together, these observations suggest that CGI promoters can accommodate elevated negative supercoiling within a spatially constrained domain, thereby functioning as local topological sinks (Fig. 3a).

**Figure 3:**
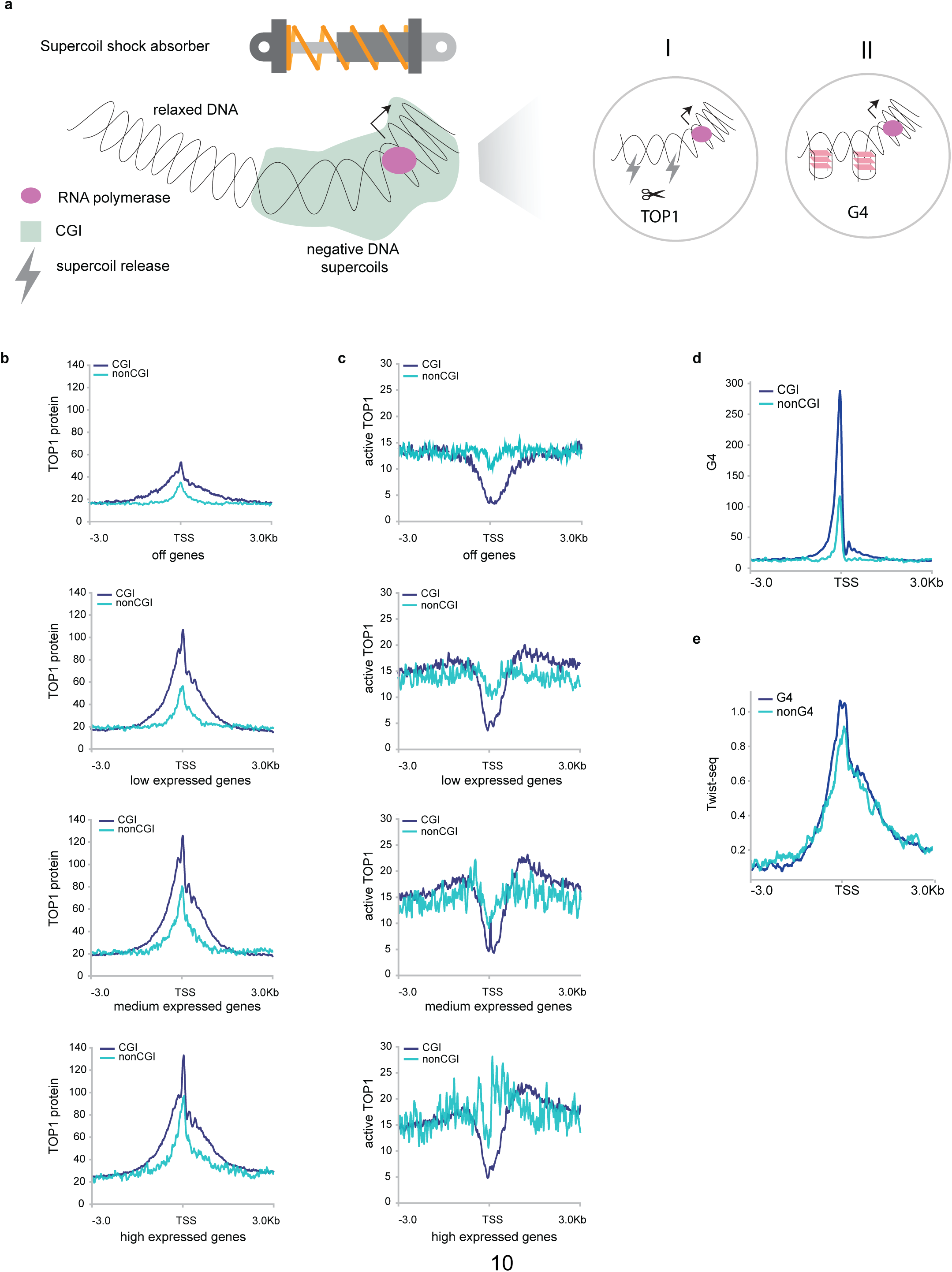
TOP1 is depleted, and G-quadruplexes are enriched, at CGIs. **a,** Model to explore how CGIs could function as negative DNA supercoil shock absorbers. **b,** TOP1-ChIP data (RPKM) (mapping TOP1 protein binding) at TSS (±3 kb) for CGI and non-CGI genes split by expression group (off, low, medium and high expressed genes)^20^. **c,** TOP1-seq data (RPKM)^20^ mapping catalytically active TOP1 at TSS (±3 kb) for CGI and non-CGI genes split by expression group (off, low, medium and high expressed genes). **d,** G4 access data mapping G4s in K562 cells showing signal enrichment in 10 bp bins across CGI and non-CGI promoters^22^. **e,** Twist-seq signal averaged across ±3 kb around TSSs of high expressing genes either containing a G4 peak within ±500 bp of the TSS (G4) or lacking such a peak (non-G4), based on G4 access peaks.

### TOP1 is depleted and G-quadruplexes are enriched at CGIs

How might CGIs function as DNA supercoil sinks? (Fig. 3a). One obvious mechanism could be differential topoisomerase activity at CGI and non-CGI promoters. TOP1 is the primary relaxase of transcription-induced DNA supercoils, associating with elongating Pol II and relieving torsional stress through transient single-strand nicking of DNA^20^. Consistent with this, TOP1 ChIP-seq revealed elevated TOP1 occupancy at CGI promoter transcription start sites (TSSs) (Fig. 3b). However, TOP1-seq, which maps catalytically engaged TOP1, showed a marked depletion of TOP1 activity at these same regions (Fig. 3c). This indicates that, despite being recruited to CGI promoters, TOP1-mediated relaxation is reduced at these sites, arguing against a primary role for TOP1 in mediating supercoil dissipation at CGIs. In contrast, we found TOP1 activity was enriched at highly expressed non-CGI promoters, suggesting that these loci rely more directly on topoisomerase-mediated relaxation to manage transcription-induced supercoiling. For a given supercoiling diffusion rate, reduced TOP1 activity would be expected to increase the characteristic propagation length of negative supercoiling (Fig. 2f). However, despite exhibiting reduced TOP1 activity (Fig. 3c), CGI promoters had supercoiling domains similar in size to those at non-CGI promoters (Fig. 2c). This suggests that CGI- specific properties may act to constrain transcription-generated torsional stress locally rather than allowing it to propagate along the surrounding chromatin.

Several mechanisms have been proposed to regulate TOP1 activity at promoters, including Pol II pausing and interactions with G-quadruplex (G4) DNA structures^21, 22^. G4s are non-canonical secondary structures formed in G-rich sequences through the stacking of guanine tetrads and are frequently associated with CpG islands. Notably, negative supercoiling generated during transcription has been proposed to promote G4 formation, which in turn can inhibit TOP1 activity and contribute to maintaining an open chromatin state at the promoter^21^. Using G4access, a genome-wide method for mapping G4 structures^22^, we confirmed that G4s are enriched at CGI promoters (Fig. 3d). However, binning highly expressed genes by G4 status revealed comparable levels of negative supercoiling regardless of G4 enrichment (Fig. 3e), indicating that G4 formation alone does not account for the elevated negative supercoiling or its localised accumulation at CGI promoters.

### Molecular dynamics simulations and ssDNA-seq reveal localised melting at CGI promoters

As bioinformatic analyses indicate that neither topoisomerase activity nor G-quadruplex formation can account for the accumulation and buffering of supercoiling at CGI promoters, we next asked whether this behaviour could arise directly from the intrinsic, sequence-dependent physical properties of DNA. To test this, coarse-grained molecular dynamics simulations using oxDNA^23,24^ were performed, which accounts for supercoiling-dependent and sequence-dependent effects in local melting and flexibility (Fig. 4a). Negatively supercoiled DNA fragments were simulated corresponding to several representative CGI and non-CGI promoter sequences under controlled torsional constraints, enabling direct comparison of their structural responses to equivalent levels of superhelical stress (Fig. 4b,c). All sequences were subjected to negative supercoil densities of σ = −0.03, −0.06 and −0.09, representing, low, intermediate and high levels of negative supercoiling respectively. An applied stretching force of 1 pN was also included, consistent with forces exerted by transcribing RNAPII^25^ (Fig. 4b,c and Fig. S3a,b; see methods).

**Figure 4:**
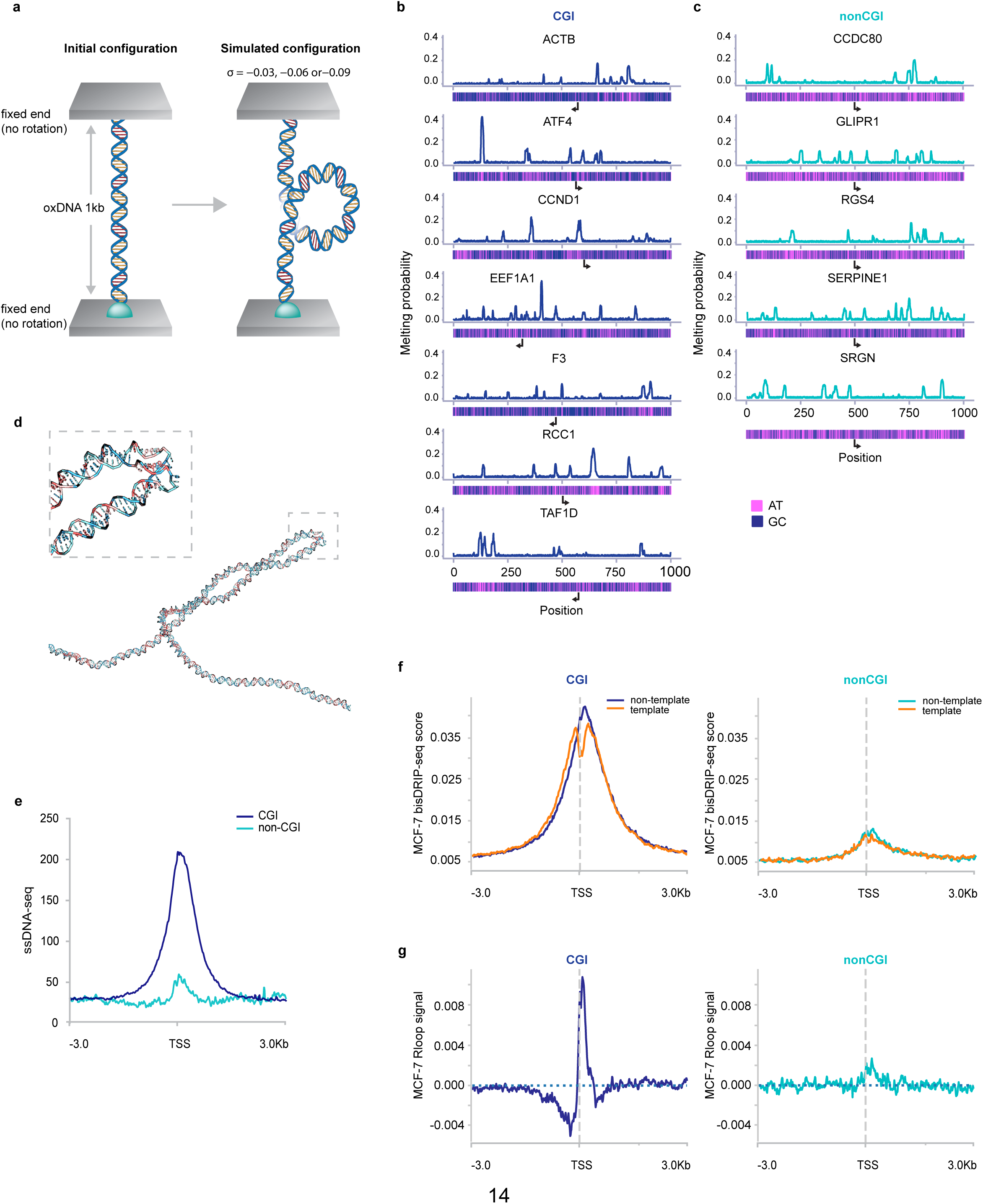
Molecular dynamics simulations and ssDNA-seq reveal localised melting at CGI promoters. **a,** OxDNA molecular dynamic simulation schematic. **b, c,** Melting profile for CGI (b) and non-CGI (c) promoter DNA sequences with a supercoil density of σ=-0.06. GC/AT content heatmaps are shown under each simulation and arrows highlight the position and direction of TSS. **d,** Snapshot of denaturation bubble formation at the tip of plectaneme for CGI gene promoter. **e,** ssDNA-seq signal^18^ (RPKM, 1 bp bins) averaged across ±3 kb around TSSs of CGI and non-CGI genes in expression group high. **f,** Mean strand-resolved bisDRIP-seq signal from MCF-7 cells^27^ is shown across CGI and non-CGI promoters centred on the TSS. Signals were oriented according to gene strand to distinguish the non-template and template DNA strands. CGI promoters show substantially greater bisDRIP signal on both strands than non-CGI promoters. **g,** Strand asymmetry of bisDRIP-seq signal at CGI and non-CGI promoters. Difference between non-template- and template-strand bisDRIP-seq signal across CGI and non-CGI promoters. Positive values indicate greater signal on the non-template strand and negative values greater signal on the template strand. CGI promoters show a localised non-template-strand bias immediately downstream of the TSS, consistent with promoter-associated R-loop formation.

Across the simulated sequences, negative superhelical densities of σ = −0.06 and −0.09 promoted the formation of denaturation bubbles, consistent with previous theoretical and experimental studies (Fig. 4d). Although melting occurred in both CGI and non-CGI sequences, the spatial distribution of these events differed significantly between promoter classes. In CGI-derived sequences, DNA melting was localised to short regions of up to approximately 20 bp, consistently occurring at discrete AT-rich pockets embedded within a more thermodynamically stable GC-rich background (Fig. 4b). In contrast, non-CGI sequences exhibited more dispersed and heterogeneous melting patterns, with multiple low-probability denaturation sites distributed throughout the sequence (Fig. 4c).

To quantify this behaviour, we measured the spatial organisation of DNA melting along each sequence using the Gini coefficient, where higher values indicate that melting is concentrated at fewer positions, and Shannon entropy, where lower values similarly indicate more localised melting (Fig. S3c; see Methods). CGI sequences displayed significantly higher Gini coefficients and lower Shannon entropy values, indicating that melting was concentrated at a smaller number of preferred sites. Conversely, non-CGI sequences exhibited lower Gini coefficients and higher entropy, reflecting a more broadly distributed pattern of melting (Fig. S3c). These results suggest that the sequence architecture of CGIs promotes spatially focused DNA melting under torsional stress, whereas melting at non-CGI promoters is more broadly distributed. These findings align with a previous *in vitro* study showing that, under low levels of negative DNA supercoiling, GC-rich DNA melts more readily than GC-poor DNA, despite GC base pairs being intrinsically more stable than AT base pairs in the absence of torsional stress^16^.

To test the simulation prediction that CGI sequence encodes the formation of localised denaturation bubbles under torsional stress, we analysed ssDNA-seq data to examine the distribution of single-stranded DNA^18^. Consistent with our hypothesis, we found that denatured ssDNA regions were significantly enriched at CGI promoters (Fig. 4e and Fig. S3d). Stratifying CGIs by activity revealed that the most active promoters exhibited the highest levels of ssDNA (Fig. 4e), consistent with the elevated negative supercoiling observed at these loci (Fig. 2c). As R-loop formation can also generate exposed ssDNA, we analysed published DRIP-seq data to examine R-loop-associated signals at gene promoters^26^. Highly expressed CGI promoters showed greater promoter-associated DRIP-seq signal than highly expressed non-CGI promoters in RPE1 cells (Fig. S3e). However, as conventional DRIP-seq provides limited spatial information and S9.6 enrichment can include non-R-loop single-stranded DNA structures, we additionally examined published strand-resolved bisDRIP-seq data from MCF-7 cells^27^. This revealed strong bisDRIP signal on both strands at CGI promoters (Fig. 4f), together with localised strand asymmetry around the TSS consistent with R-loop formation (Fig. 4g). Thus, R-loops are likely to contribute to strand exposure at active CGI promoters but occur within a broader region of CGI-associated ssDNA. Together, these observations support the presence increased DNA strand exposure at active CGI promoters, consistent with the dynamic, localised supercoiling-induced DNA melting predicted by the oxDNA simulations.

However, whilst CGI and non-CGI sequences exhibited distinct spatial patterns of supercoiling-induced DNA melting (Fig. 4b,c), the oxDNA simulations predicted relatively modest differences in overall melting under equivalent torsional stress. In contrast, the *in vivo* data revealed a much more pronounced difference in DNA strand exposure, with highly expressed non-CGI promoters exhibiting substantially less ssDNA and R-loop-associated signal than CGI promoters, despite showing detectable negative supercoiling (Fig. 2c). Thus, although intrinsic sequence properties influence where supercoiling-induced melting occurs, they cannot fully account for the pronounced difference in strand exposure observed between CGI and non-CGI promoters in vivo. This suggests that additional features of the promoter environment contribute to the elevated ssDNA observed at active CGI promoters.

### CGI promoters act as topological sinks through localised DNA melting and writhe at nucleosome-depleted regions

One prominent difference between CGI and non-CGI promoters is their nucleosome organisation, with CGI promoters characterised by more pronounced nucleosome depletion around the TSS^15^. We therefore asked whether this difference could contribute to their distinct structural responses to torsional stress. Analysis of Fiber-seq–derived nucleosome occupancy maps^28,29^ confirmed distinct nucleosome occupancy profiles at CGI and non-CGI promoters (Fig. S3f, g). This difference could arise, at least in part, from intrinsic sequence properties, as previous *in vitro* studies have shown that the sequence composition of CGI DNA intrinsically disfavors nucleosome formation^15^. Consistent with this, sequence-based analysis using DNAcycP2^30^, a model trained on experimental loop-seq data^31,32^ to predict intrinsic DNA cyclizability, revealed a substantial decrease in predicted cyclizability around CGI TSSs, with no comparable decrease at non-CGI promoters (Fig. S3h).

We therefore first incorporated sequence-dependent nucleosome occupancy into the oxDNA simulations (Fig. S4a) and then extended the model by imposing mechanical constraints on nucleosome-bound DNA, reflecting the restricted conformational freedom associated with DNA wrapping around histone octamers (Fig. 5a,b; see Methods). Incorporating nucleosome occupancy alone preferentially confined melting to nucleosome-free regions (Fig. S4a). However, the pronounced difference in strand separation between CGI and non-CGI promoters observed experimentally by ssDNA-seq only emerged in the simulations when these mechanical constraints were imposed (Fig. 5a,b). Under these conditions, CGI promoters retained nucleosome-depleted regions around the TSS that supported localised melting under negative supercoiling (Fig. 5a,c,d and Fig. S4b). In contrast, higher nucleosome occupancy around non-CGI TSSs restricted promoter-proximal melting, with many AT-rich, melting-prone regions located within mechanically constrained nucleosome-bound DNA, leaving fragmented sites of denaturation in linker DNA distal to the promoter (Fig. 5b,c). Thus, incorporating nucleosome organisation and its mechanical constraints amplified the differences in melting behaviour observed in the unconstrained simulations, producing focused promoter-proximal melting at CGI promoters but infrequent, dispersed melting at non-CGI promoters.

**Figure 5:**
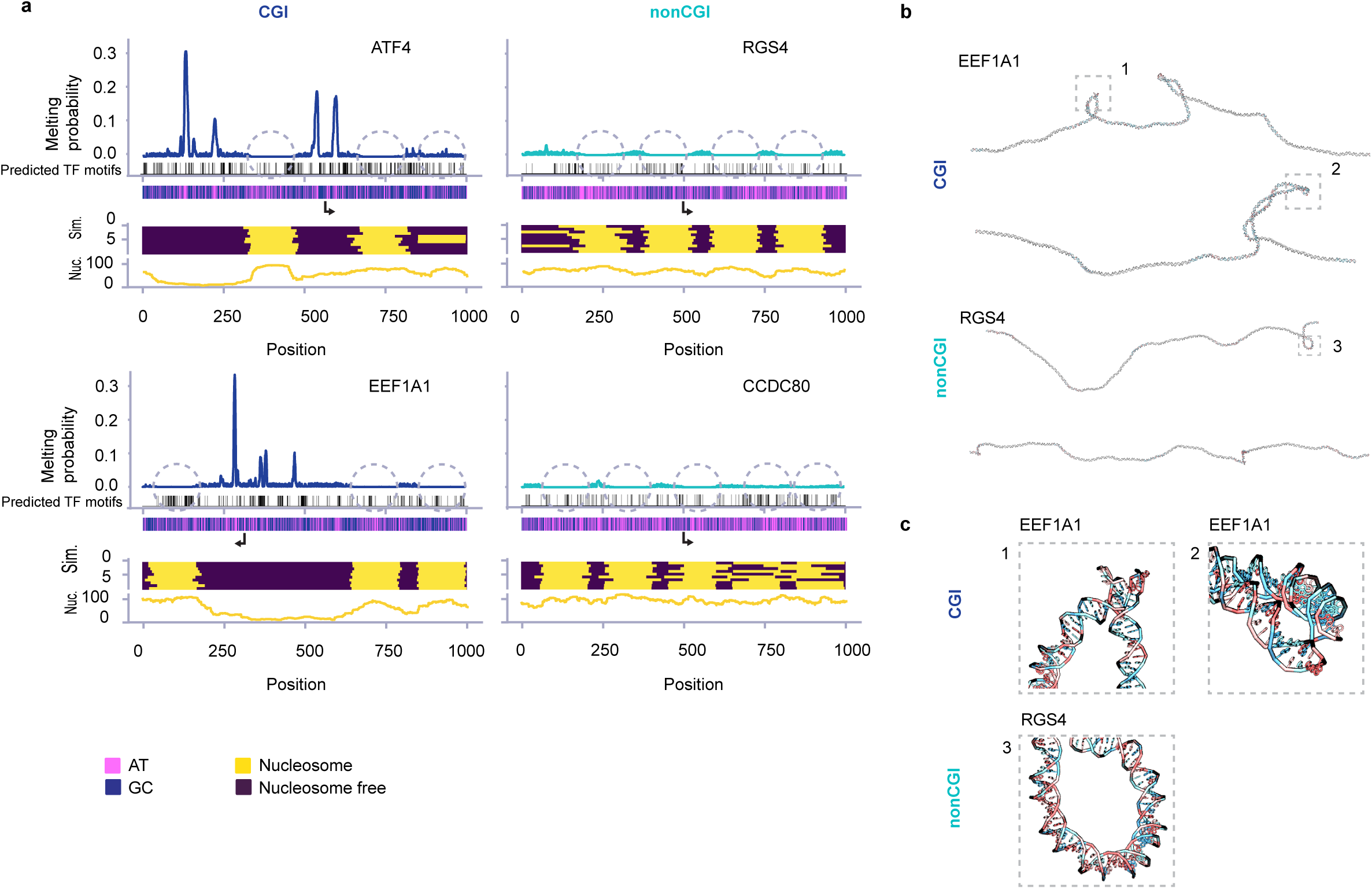
CGI promoters act as topological sinks through localised DNA melting and writhe at nucleosome-depleted regions. **a,b,** Melting profile for CGI (a) and non-CGI (b) promoter DNA sequences with supercoil density of σ=-0.06, with nucleosome-bound regions (dashed circles) modelled as mechanical constraints that restrict local DNA melting, bending and twisting. Nucleosome positioning was predicted from Monte Carlo based simulations (Sim track) of fibre seq data (Nuc track). Vertical ticks beneath each profile indicate the positions of predicted TF motif occurrences identified by FIMO, irrespective of TF identity. All panels are shown on the same y-axis scale. **c,** Simulation snapshots illustrating DNA writhe and melting at CGI promoters. Dashed boxes indicate regions of local DNA melting shown at higher magnification in **d,** Magnified views of the boxed regions in c, illustrating local DNA melting at AT-rich sequence pockets. **e,** Somatic mutation rates at CGI and non-CGI promoters across expression groups. Forest plot showing CGI/non-CGI somatic mutation rate ratios within ±500 bp of the TSS for transcriptionally inactive (Off), low-, medium- and highly expressed promoters. Points indicate estimated rate ratios and horizontal lines indicate 95% confidence intervals from negative-binomial mixed-effects models accounting for promoter sequence opportunity, tumour-wide mutation burden and tumour-level heterogeneity. The dashed line at rate ratio = 1 indicates no difference between CGI and non-CGI promoters. The relationship between CGI status and mutation rate differed significantly across expression groups (CGI status × expression interaction, likelihood-ratio test, P = 4.8 × 10⁻⁷).

Nucleosome depletion could additionally contribute to the distinctive Twist-seq profile at CGI promoters by altering how local torsional stress is partitioned. Nucleosome formation constrains approximately one negative unit of linking number^33^, and nucleosome-depleted DNA is therefore expected to accommodate torsional stress differently from nucleosome-occupied DNA. In nucleosome-free DNA, negative torsional stress can be accommodated through both DNA undertwist and writhe. The undertwist component would favour bTMP intercalation and could therefore contribute to the experimentally observed Twist-seq signal. Our simulations further predict that negative torsional stress within CGI nucleosome-depleted regions is also accommodated through local writhe, accompanied by focused melting of 10– 20 bp AT-rich pockets at or near plectonemic apices (Fig. 5c,d and Fig. S4b). Consistent with this structural response, CGI promoter sequences exhibited greater DNA shortening than non-CGI sequences under the same applied stretching force of 1 pN (Fig. S4c). This coupling of local melting and writhe is consistent with previous studies of supercoiled naked DNA^34,35^. Additional analysis revealed pronounced, localised changes in internal energy associated with DNA melting at CGI promoters, contrasting with the smaller changes observed at non-CGI promoters (Fig, S5 and Fig. S6). Thus, whereas the reaction–diffusion model provides a physical framework for the spatial extent of supercoiling at CGI and non-CGI promoters, the oxDNA simulations reveal how sequence-dependent nucleosome organisation and DNA mechanics could generate the distinctive promoter-proximal supercoiling and structural behaviour observed at CGIs.

Because local DNA melting could disrupt the duplex structure required for transcription factor (TF) binding, we asked whether supercoiling-induced melting preferentially occurred outside predicted TF-binding motifs (Fig. 5a,b and Fig. S7a)^36,37^. Across the 12 nucleosome-positioned promoter simulations, predicted TF motif sequences consistently exhibited lower melting probabilities than their immediately flanking DNA (Fig. S7b). In the final nucleosome- and topologically constrained simulations, the two CGI promoters that developed high-probability melting regions, ATF4 and EEF1A1, showed significantly fewer overlaps between TF motifs and these regions than expected from within-promoter positional randomisation (Fig. S7c). Thus, supercoiling-induced melting was spatially organised within the simulated promoter sequences, with the strongest melting preferentially occurring outside predicted TF motifs.

Given the increased propensity of highly transcribed CGI promoters to undergo strand separation, we next asked whether this was associated with increased somatic TES frequency. Analysis of somatic mutations from the Pan-Cancer Analysis of Whole Genomes (PCAWG) dataset^38^ revealed an expression-dependent relationship between promoter class and mutation rate (interaction P = 4.8 × 10⁻⁷), with CGI promoters showing a lower relative mutation rate in transcriptionally inactive genes but a higher relative mutation rate among highly expressed genes (Fig. 5e). Among highly expressed genes, CGI promoters showed a 10.6% higher mutation rate within ±500 bp of the TSS than non-CGI promoters (rate ratio = 1.106, 95% CI = 1.026–1.192, P = 0.0083). This raises the possibility that increased exposure of ssDNA associated with transcription-induced torsional stress may contribute to the elevated mutation frequency observed at highly expressed CGI promoters.

Together, these findings support a model in which the intrinsic sequence properties and characteristic nucleosome organisation of CGI promoters enable them to act as local sinks for transcription-induced torsional stress (Fig. 6). CGI sequence intrinsically disfavours nucleosome formation^15^ and exhibits reduced predicted DNA cyclizability^31,32,39^, contributing to nucleosome depletion around the TSS. This provides a local environment in which transcription-generated negative torsional stress can be accommodated through coupled DNA undertwist, writhe and melting, whereas greater nucleosome occupancy at non-CGI promoters restricts these structural responses. We propose that this capacity for local structural accommodation enables CGI promoters to function as topological sinks.

**Figure 6:**
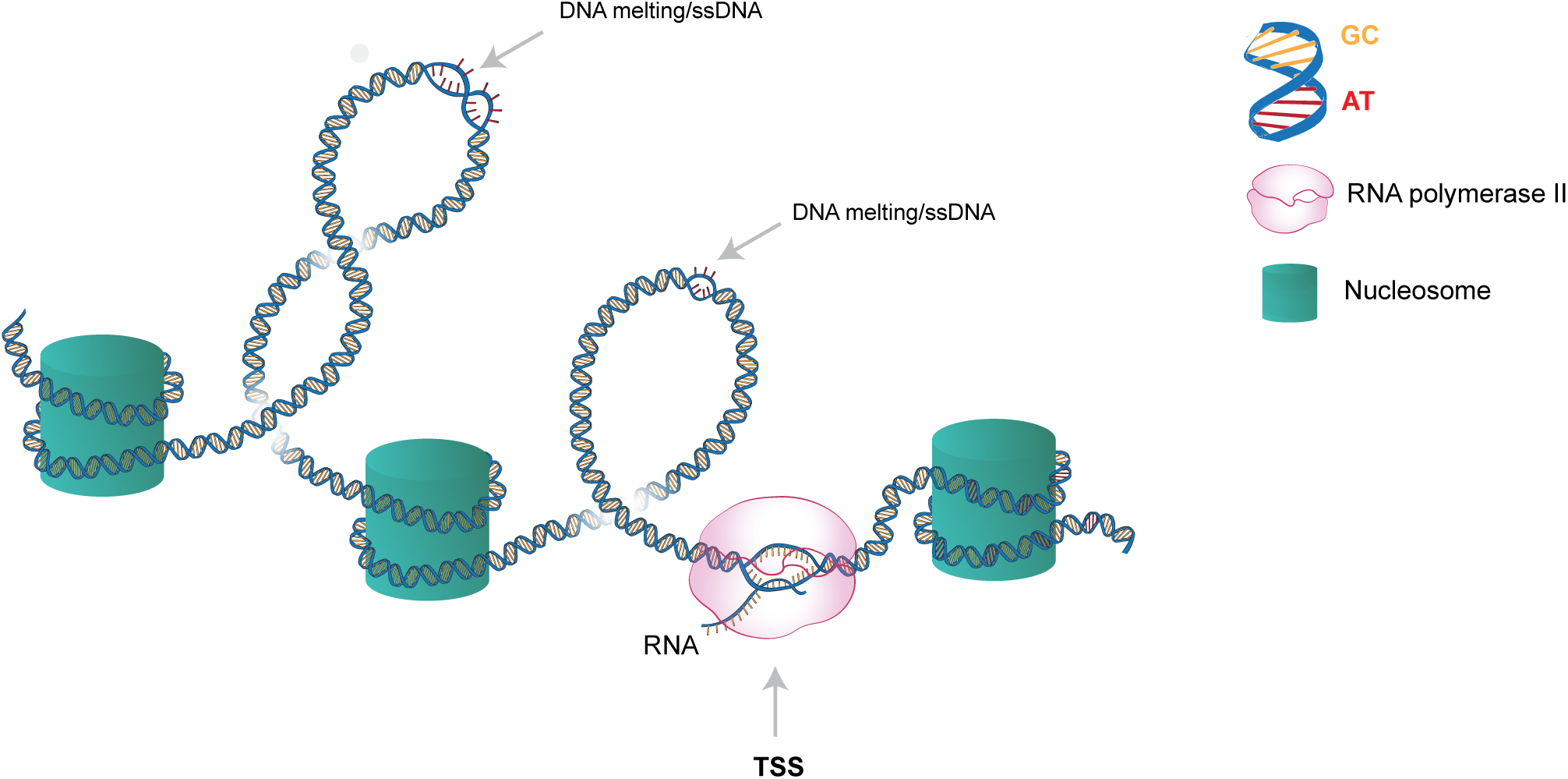
Model for CpG island promoters as topological sinks for transcription-generated supercoiling. The distinctive sequence (GC rich (yellow base pairs)) and chromatin architecture of CGI promoters create a nucleosome-depleted region in which transcription-generated torsional stress can be accommodated through DNA underwinding, writhe and local DNA melting. RNA polymerase II represents the source of transcription-generated torsional stress. Within nucleosome-free DNA, negative supercoiling promotes plectoneme formation, with localised melting preferentially occurring at embedded AT-rich pockets (red base pairs) near plectoneme tips. This partitioning provides a mechanism by which CGI promoters can buffer and redistribute transcription-generated torsional stress, while topoisomerase activity provides a route for its relaxation

To examine how CGI promoters accommodate increased transcription-generated torsional stress, we measured DNA supercoiling following acute heat-shock-induced transcriptional activation. Heat-shock-induced genes were identified using previously generated TT-seq data (Supplementary Table 3), and their transcriptional response was further examined using TT-seq data generated under our experimental treatment conditions (Supplementary Fig. 8a). CGI promoter Twist-seq signal showed no significant change following heat shock (Fig. 7b,c; P = 0.071). However, TOP2 inhibition during heat shock increased CGI promoter Twist-seq signal (Fig. 7b,c), consistent with a role for TOP2 in relaxing transcription-generated torsional stress, potentially through the resolution of DNA writhe. In contrast, and consistent with TOP1-seq data, TOP1 inhibition had no significant effect on CGI promoter Twist-seq signal. Importantly, the increase in CGI promoter Twist-seq signal following TOP2 inhibition persisted after matching CGI and non-CGI genes for expression (Fig. 7d,e, Fig. S8b and Supplementary Table 4), suggesting that this response is not simply attributable to differences in transcriptional output between the two promoter classes. Together, these results support a model in which CGI promoters accommodate transcription-induced torsional stress through the combined effects of intrinsic DNA properties, nucleosome organisation and topoisomerase-mediated relaxation, contributing to their proposed function as topological sinks.

**Figure 7:**
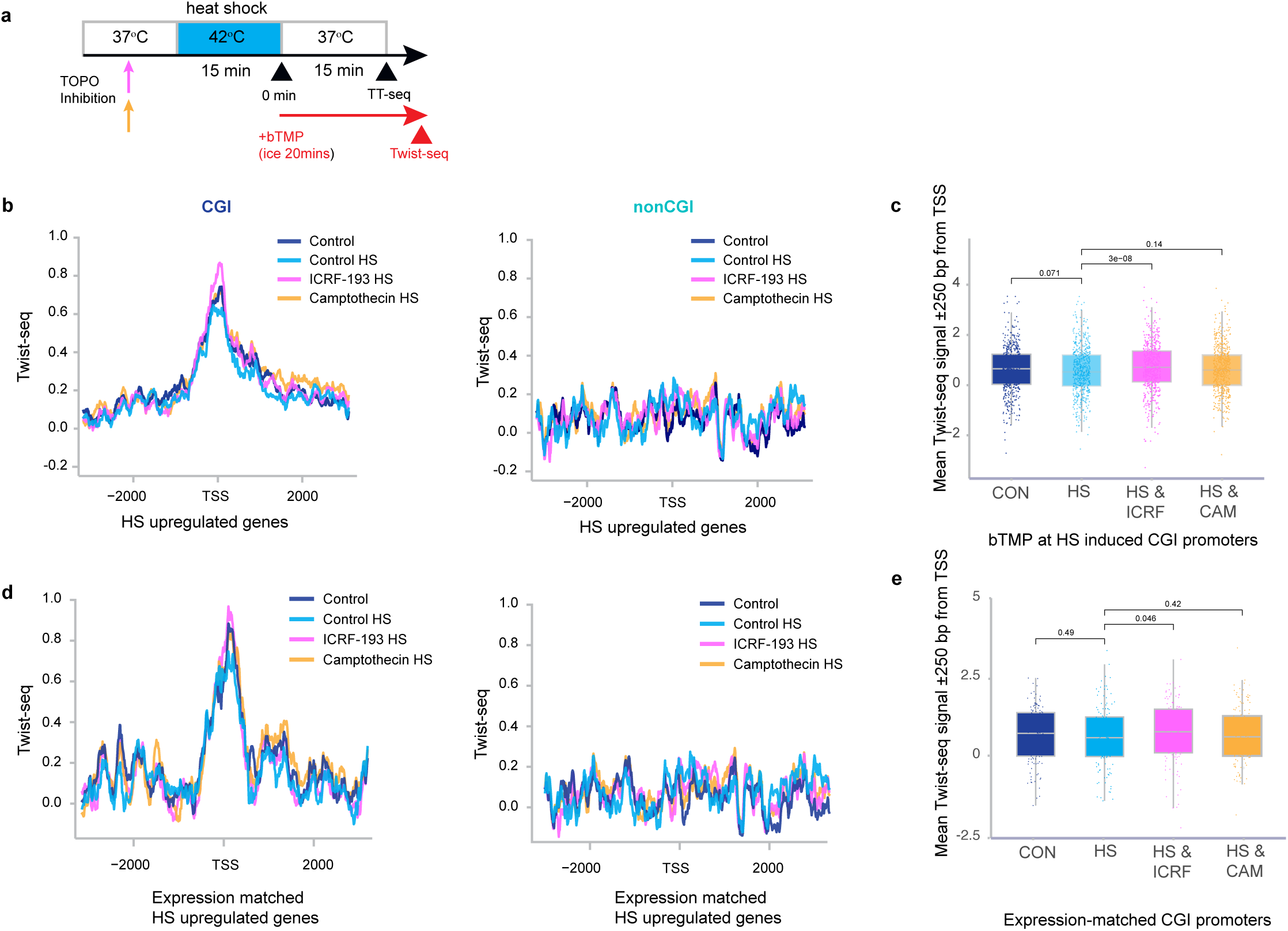
TOP2 inhibition increases Twist-seq signal at CGI promoters during transcriptional activation. **a,** Schematic of the experimental design showing topoisomerase inhibitor pretreatment (3 h), heat shock (42 °C, 15 min) and subsequent bTMP labelling (20 min on ice) for Twist-seq, or recovery at 37 °C (15 min) followed by TT-seq labelling. **b,**Twist-seq signal profiles (1-bp bins) ±3 kb around the transcription start sites (TSSs) of heat-shock-induced CGI (left) and non-CGI (right) genes (TT-seq log₂ fold change > 1, adjusted P < 0.05) under control (CON), heat shock (HS), TOP2 inhibition during heat shock (HS & ICRF-193 (ICRF)) and TOP1 inhibition during heat shock (HS & camptothecin (CAM)) conditions (n = 554 CGI and n = 143 non-CGI promoters; Supplementary Table 3). **c,** Boxplots showing mean Twist-seq signal within ±250 bp of the TSS for CGI promoters of heat-shock-induced genes under the conditions described in b (n = 554). Individual points represent promoters; boxes indicate the median and interquartile range. Statistical significance was determined using paired two-sided Wilcoxon signed-rank tests (unadjusted P values). CON versus HS, P = 0.071; HS versus HS & ICRF, P = 3.0 × 10⁻⁸; HS versus HS & CAM, P = 0.144. **d,** Twist-seq signal profiles (1-bp bins) ±3 kb around the TSSs of expression-matched heat-shock-induced CGI (left) and non-CGI (right) promoters under the conditions described in b. CGI and non-CGI promoters were matched according to their transcriptional output following heat shock, measured by TT-seq (n = 129 promoters per group; Supplementary Table 4). **e,** Boxplots showing mean Twist-seq signal within ±250 bp of the TSS for expression-matched CGI promoters under the conditions described in b (n = 129). Individual points represent promoters; boxes indicate the median and interquartile range. Statistical significance was determined using paired two-sided Wilcoxon signed-rank tests (unadjusted P values). CON versus HS, P = 0.49; HS versus HS & ICRF, P = 0.046; HS versus HS & CAM, P = 0.42.

## Discussion

Our results reveal a sequence-encoded mechanism by which CGIs regulate DNA supercoiling and transcription at gene promoters. By integrating genome-wide supercoiling maps with molecular dynamics simulations, reaction-diffusion modelling and genomic analyses, we show that the distinctive sequence and chromatin architecture of CGI promoters enable transcription-induced torsional stress to be accommodated through DNA underwinding, writhe and localised melting at embedded AT-rich pockets. CGI promoters may therefore function as topological sinks, or ‘shock absorbers’, that buffer transcription-induced torsional stress, while the resulting promoter topology could favour polymerase engagement and transcriptional initiation (Fig. 6).

The spatial distribution of supercoil-induced DNA melting in our simulations further emphasises the importance of local chromatin architecture. Melting occurred preferentially within nucleosome-free regions and was largely distinct from predicted transcription factor binding motifs. Recent work has shown that nucleosome positioning influences transcription factor access to DNA in a factor-specific manner ^40^. Conversely, transcription factor binding can establish defined translational and rotational positioning of adjacent nucleosomes^41^, revealing an organised relationship between regulatory factor occupancy and local chromatin architecture. This suggests that nucleosome organisation and local DNA sequence jointly influence where negative torsional stress is accommodated through strand separation, with nucleosome-free and linker DNA providing preferential sites for supercoil-induced melting.

These findings provide a new perspective on the functional importance of CGI sequence composition. CGIs have long been associated with transcriptional regulation through their roles in promoting open chromatin states, guiding histone modifications and resisting DNA methylation. Studies demonstrating that synthetic CGI-like sequences can establish promoter-like chromatin states^42^, and that CGI sequences remain hypomethylated even when relocated across species^43^, highlight the extent to which these properties are encoded by DNA sequence itself. Our results suggest that CGI sequence composition may additionally encode intrinsic mechanical properties that influence how promoter DNA responds to transcription-generated torsional stress. This idea is supported by previous work demonstrating that GC-rich DNA can melt more readily than GC-poor DNA under low levels of negative supercoiling, despite the greater intrinsic stability of GC base pairs^16^ and by atomistic simulations showing that negative supercoiling promotes denaturation at AT-rich sequences, potentially influencing plectoneme positioning^35^. More broadly, transcription-generated torsional stress has been shown to promote sequence-dependent structural transitions, including R-loop formation and G-quadruplex stabilisation^44^, while DNA-embedded ribonucleotides enriched at CpG islands have been implicated in modulating transcription-associated negative supercoiling through RNase H2-mediated processing^45^. The capacity to accommodate torsional stress also extends to chromatin, where single-molecule experiments have demonstrated that changes in nucleosomal entry–exit DNA geometry can buffer transcription-generated positive torsional stress and facilitate Pol II progression through nucleosomes^46^. Together, these findings illustrate how DNA sequence and chromatin architecture influence the response to transcription-generated torsional stress, while our results suggest that CGI promoters provide a specialised environment for accommodating negative supercoiling.

An important implication of this model concerns CpG methylation, which occurs at CGIs during ageing and disease and is usually associated with lower transcriptional activity^47^. A recent study demonstrated that recognition of unmethylated CpGs by the DNMT1 CXXC domain inhibits DNA methyltransferase activity, providing a mechanism that protects CGIs from inappropriate methylation^48^. Cytosine methylation at CGIs has also been shown to affect the biophysical properties of DNA, modulating its interaction with nucleosomes in a context dependent manner, while methylation is generally thought to stabilise nucleosome formation^49^. Importantly, cytosine methylation also stabilises the DNA duplex against melting, increasing its melting temperature^50^, suggesting that aberrant CGI methylation may alter the susceptibility of promoter DNA to supercoil-induced melting and consequently its capacity to accommodate transcription-generated torsional stress, with potential consequences for transcriptional regulation.

Comparative genomic analyses have shown that CGI promoters evolve more slowly than non-CGI promoters^51^ and are associated with genes under strong functional constraint^52^. The GC-rich sequence environment of CGIs, which favours nucleosome depletion, together with locally positioned AT-rich regions susceptible to supercoil-induced melting, may confer mechanical properties that support sustained transcription under high torsional load. We propose that the capacity to buffer and redistribute negative supercoiling through localised DNA structural transitions may represent an additional, selective pressure acting on CGI promoter sequences. The expression-dependent relationship between CGI status and somatic mutation rate further highlights the potential consequences of transcription-associated torsional stress for genome stability, as transient strand exposure may create opportunities for DNA damage and mutagenesis. Thus, the capacity of CGI promoters to accommodate torsional stress may involve a balance between maintaining topological homeostasis during sustained transcription and limiting the potential genomic consequences of transient DNA strand exposure.

## Materials and Methods

### Cell culture

RPE1 cells (ATCC CRL-4000) were cultured in DMEM-F12, 3 mM glutamine and 15 mM HEPES supplemented with 0.34% sodium bicarbonate, 10% FBS, penicillin (100 U ml−1), streptomycin (100 μg ml−1) and phenol red (8.1 mg l−1).

### Twist-seq

RPE1 cells or control genomic DNA were treated with 500 μg/ml of bTMP for 20 mins on ice in the dark. bTMP was UV cross-linked to DNA at 360 nm for 10 min. DNA was purified from cells using SDS and proteinase K digestion followed by phenol-chloroform-isoamyl alcohol extraction. DNA was fragmented by sonication (13 times for 30s on/off Biorupter Pico). Biotin incorporation into DNA was detected by dot blotting using alkaline phosphatase−conjugated avidin as a probe. The bTMP−DNA complex in TE was immunoprecipitated using avidin conjugated to magnetic beads for 2 h at room temperature and then overnight at 4 °C. Beads were washed sequentially for 5 min each at room temperature with TSE I (20 mM Tris, pH 8.1, 2 mM EDTA, 150 mM NaCl, 1% Triton X-100 and 0.1% SDS), TSE II (20 mM Tris, pH 8.1, 2 mM EDTA, 500 mM NaCl, 1% Triton X-100 and 0.1% SDS) and buffer III (10 mM Tris, pH 8.1, 0.25 M LiCl, 1 mM EDTA, 1% NP40 and 1% deoxycholate). Beads were then washed twice with TE buffer for 5 min. To extract the bTMP-bound DNA the samples were boiled for 10 min at 90 °C in 50 μl of 95% formamide with 10 mM EDTA. Samples were then made up to 400 μl with water, and the bTMP-bound ssDNA was precipitated by EtOH precipitation on dry ice for a minimum of 30 min or −80 °C overnight. DNA was eluted in 15 μl low TE and the full amount used to prepare sequencing libraries with xGen™ ssDNA & Low-Input DNA Library Preparation Kit (IDT 10009859) following the manufacturer’s instructions. Library PCR amplification was typically 16 cycles after which libraries were sized and quality controlled on a D1000 Tapestation tape (Agilent). Illumina sequencing (NovaSeq X Plus Series (PE150; paired-end DNA-seq of 150-bp read length) to give a read depth of at least 200M reads per sample. FASTQ sequence files were obtained and the Twist-seq reads were adapter- and quality-trimmed using TrimGalore v0.6.6 and aligned to the Human (hg38) reference genome using Bowtie2 v2.5.3. Aligned reads were processed with Samtools v1.6, and the deepTools “bamCoverage” tool with RPKM normalization, bin size of 1bp and with blacklisted regions removed. Sequence read depth for RPE1 control replicates were 215 million, 269 million, 575 million and 452 million reads respectively with 83%, 91%, 93% and 94% mapping efficiency. To release torsional stress in the chromatin fiber, the DNA was cleaved by treatment of cells with 100 μM bleomycin (Sigma B2434) for 10 min at 37 °C prior to bTMP treatment. Sequence read depth for the bleomycin treated replicates were 681 and 275 million reads respectively with 96% mapping efficiency. For topoisimerase inhibition, RPE1 cells were pretreated with ICRF-193 (35 μM, Enzo BML-GR332-0001) or camptothecin (5 μM, Sigma C9911) for 3 h before heat shock at 42 °C for 15 min. Cells were subsequently labelled with bTMP for 20 min prior to Twist-seq.

### TT-seq

Nascent RNA was labelled by adding 500 µM 4-thiouridine (4sU)(Sigma, T4509) to RPE1 cells in T75 flasks and incubating at 37 °C for 10 min. Media was aspirated and RNA extraction was performed with TRIzol (Invitrogen) following the manufacturers’ instructions. After DNase treatment (Turbo DNase, Thermo Fisher Scientific) RNA concentration and purity were determined using a NanoDrop. RNA (70 µg) was fragmented in 100 µl H20 to <1.5 kb by 20 cycles of 30 s on/30 s off at high power in a Biorupter plus and RNA size assessed by agarose gel electrophoresis. Fragmented 4sU labelled RNA was biotinylated by adding 140 µl of EZ-Link Biotin-HPDP (1 mg.ml−1 in dimethylformamide; Pierce, 21341), 70 µl of 10x biotinylation buffer (100 mM Tris-HCl pH 7.5, 10 mM EDTA) and H20 to a final volume of 700 µl. This was incubated at room temperature for 1.5 h with rotation. Unincorporated biotin-HPDP was removed by two rounds of chloroform extraction with 2 ml Phase lock gel heavy tubes (Eppendorf). RNA was precipitated with 1/10 volume of 5 M NaCl and an equal volume of Isopropanol. This was inverted to mix and incubated at room temperature for 10 min followed by centrifugation at 10,000 g for 20 min at room temperature. RNA pellet was washed with 80% EtOH and centrifuged at 13,000 rpm (15871 g) for 10 min at 4 °C. RNA was resuspended in 100 µl H20 and dissolved by heating to 40 °C for 10 min with agitation. RNA was then immediately placed on ice and RNA concentration determined by Nanodrop spectrophotometer. Biotinylated 4sU labelled RNAs were then recovered using µMACS Streptavidin MicroBeads (Miltenyi, 130-074-101) and separation on a µMACS Separator. For the concentration of total RNA in µg per sample an equal amount in µl of Streptavidin microbeads was added. This was incubated at room temperature for 15 min with rotation. µMacs columns were equilibrated with 900 µl room temperature washing buffer (100 mM Tris-HCl pH 7.5, 10 mM EDTA, 1 M NaCl, 0.1% Tween20). The RNA/streptavidin bead solution was then applied to the column followed by three washes with 900 µl of washing buffer at 65 °C and three washes with 900 µl of washing buffer at room temperature. RNA was eluted with 2 × 100 µl of fresh elution buffer (100 mM dithiothreitol in RNase-free H20) directly into 2 ml lobind tubes (Eppendorf) containing 700 µl Buffer RLT (RNeasy MinElute Cleanup Kit, Qiagen). 500 µl of 100% ethanol was added to the RNA solution, and mixed thoroughly by pipetting before RNA was purified through RNAeasy MinElute Spin Columns. RNA concentration was determined using a Nanodrop and libraries for RNA-seq were prepared and indexed using NEBNext® Ultra™ II Directional RNA Library Prep Kit for Illumina® (NEB #E7645L) and NEBNext Singleplex Oligos for Illumina (NEB #E7335, E7500) following the manufacturer’s instructions. Libraries were sized and quality controlled on a D1000 Tapestation tape (Agilent) and Illumina sequencing (paired-end RNA-seq of 150-bp read length) was performed on NovaSeq (Novogene, Cambridge). FASTQ sequence files were obtained and the RNA-seq reads were aligned to the Human (hg38) reference genome using Bowtie2 v2.5.3. Aligned reads were processed with Samtools v1.6, and the deepTools “bamCoverage” tool with RPKM normalization, bin size of 20bp and with blacklisted regions removed. Sequence read depth for RPE1 control replicates 146 million and 71 million reads with 86% mapping efficiency.

### Identification and analysis of heat-shock-induced genes

Heat-shock-induced genes were identified using TT-seq data generated in RPE1 cells under equivalent heat-shock conditions. Genes exhibiting a log₂ fold change > 1 and an adjusted P value < 0.05 were classified as heat-shock-induced. This identified 713 genes, comprising 554 were classified as CGI-associated and 143 as non-CGI, with 16 genes lacking CGI classification (Supplementary Table 3). The CGI and non-CGI promoter sets were used to examine Twist-seq signal following heat shock and topoisomerase inhibition.

Gene expression under these treatment conditions was independently assessed using TT-seq data generated in this study. Expression analyses were restricted to genes represented in this dataset, comprising 523 CGI and 132 non-CGI genes.

To control for differences in transcriptional output between CGI and non-CGI promoters, heat-shock-induced genes were matched according to their expression levels measured by TT-seq following heat shock. Non-CGI genes were ordered by increasing log-transformed expression (log₁₀(RPKM + 0.01)) and matched individually to the CGI gene with the closest log-transformed expression value. Matching was performed without replacement, such that each CGI gene could be selected only once. This yielded 129 expression-matched CGI and 129 non-CGI genes, which were used for subsequent Twist-seq analyses (Fig. 7d,e). The matched gene sets and their expression values are provided in Supplementary Table X.

For statistical comparisons of Twist-seq signal between treatment conditions, mean signal was calculated within ±250 bp of the TSS for each promoter. Differences between conditions were assessed using paired two-sided Wilcoxon signed-rank tests, with promoters paired across treatment conditions (Fig. 7c,e). Gene expression was quantified as RPKM and transformed as log₁₀(RPKM + 0.01). Differences in expression between treatment conditions were assessed using paired two-sided Wilcoxon signed-rank tests, with genes paired across conditions and P values adjusted for multiple comparisons using the Benjamini–Hochberg method (Supplementary Fig. 8a). Expression differences between matched CGI and non-CGI gene sets were assessed using a paired two-sided Wilcoxon signed-rank test (Supplementary Fig. 8b).

### 1D reaction-diffusion model for supercoiling dynamics near the TSS

The size of the negatively supercoiled domains around the TSS can be more quantitatively understood by modelling the dynamics of supercoiling via a reaction-diffusion equation^53^,

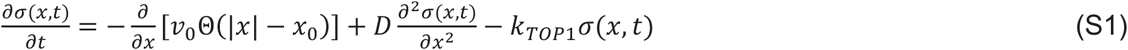

where *σ*(*x*, *t*) is the supercoiling density at position *x* and time *t*, *v*_0_ is the velocity of RNA polymerase, which is assumed to be localised betweens position ±*x*_0_, simulating the length of a typical gene with the TSS at *x* = −*x*_0_, and Θ is the Heaviside function (equal to 1 for |*x*| < *x*_0_, and 0 otherwise).

Additionally, *D* is the supercoiling diffusion coefficient, and *k_T_*_0*P*1_is the rate of supercoil relaxation by TOP1. A steady state solution of this equation for large (positive or negative) *x* is given by an exponentially decaying supercoiling density; the characteristic size of the exponential decay can then be interpreted as the supercoiling domain size size 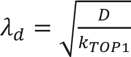. If we use the reasonable estimates of *D* ≃ 1 kbp^2^/s for the supercoiling diffusion, relevant for chromatin^53^ or writhed DNA^54^, and of *k_T_*_0*P*1_ ≃ 0.1 − 1 s^−1^ ^33^ for the TOP1 relaxation rate, the resulting domain has size *λ_d_* ≃ 1 − 3 kbp, in line with our experimental observation in Fig. 2c. Eq. (S1) is a generalisation of the model described in^53^: instead of modelling transcription as a delta-like flux centred at the instantaneous polymerase position, here we model it in an effective way, spreading it over the size of the gene being transcribed.

The steady state solution of Eq. (S1) is given by:

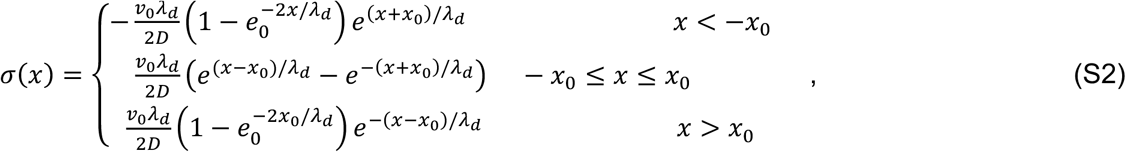

and the negative part (modelling bTMP profiles) is shown in Fig. 2f for the parameters detailed in the corresponding caption.

### oxDNA simulations

To investigate the sequence-dependent response of promoter DNA to torsional stress, we performed coarse-grained molecular dynamics simulations using the sequence-dependent oxDNA2 model^23,24^. We selected 12 representative human promoter sequences, comprising 7 CpG island (CGI) promoters and 5 non-CGI promoters. Promoters were selected from CGI and non-CGI classes based on their strong supercoiling signal in the experimental data, with the selected genes predominantly belonging to the high-expression category. For each promoter, a 1 kb DNA fragment centred on the transcription start site (TSS) was extracted and converted into the corresponding oxDNA representation. Specifically, the following promoters were chosen: CGI: ACTB, ATF4, CCND1, EEF1A1, F3, RCC1 and TAF1D, nonCGI: CCDC80, GLIPR1, RGS4, SERPINE1 and SRGN.

In oxDNA2, each nucleotide is represented as a rigid body interacting through effective potentials describing hydrogen bonding between complementary bases, stacking and cross-stacking interactions, excluded-volume effects, and backbone connectivity. The model accurately reproduces the thermodynamic, structural and mechanical properties of DNA, including flexibility, melting behaviour and supercoiling-induced structural transitions. Simulations were performed using the GPU implementation of the stand-alone oxDNA molecular dynamics code. DNA molecules were modelled as linear fragments subject to an external stretching force of 1 pN, chosen to approximate the tension experienced by DNA during active transcription. The two DNA ends were spatially constrained by repulsive boundary walls and rotationally constrained to prevent relaxation of torsional stress through end rotation. Supercoiling was imposed by fixing the total linking number of the molecule to generate superhelical densities of σ = −0.03, −0.06 and −0.09, corresponding respectively to weak, intermediate and strong levels of transcription-associated torsional stress.

All simulations were initiated from fully base-paired stretched DNA conformations. Molecular dynamics trajectories were evolved for 10^8^ integration steps. Following an initial equilibration period, which was discarded from subsequent analysis, approximately 1000 independent configurations were sampled from each trajectory for quantitative analysis. Because denaturation bubbles often persist for timescales comparable to the simulation duration, multiple independent simulations were required to adequately sample the ensemble of accessible conformations. Unless otherwise stated, 20 independent simulations were performed for each sequence and supercoiling condition using distinct initial velocity distributions. To map oxDNA timescale to real time, we consider the time needed for 1 kbp of DNA to diffuse its own radius. This can be computed to be of the order of 0.2 ms in water, or 2 ms in the cytosol; this value is computed assuming 10-fold higher viscosity in the nucleoplasm than in water, a water viscosity of 1 cP, and a diffusion coefficient of 1 kbp of 3.4 micron^2/s ^55^ (Lukacs et al., 2000). In oxDNA this is matched for a timestep of 1.5 ps or 15 ps in water or the nucleoplasm. This corresponds to a total timescale in the kymographs simulated of approximately 1.5 ms in vivo.

#### DNA melting

Base-pair melting was quantified from the disruption of native Watson-Crick base pairing. A base pair was classified as melted when the absolute value of the hydrogen-bonding energy drops below 20% of the maximum, equivalent to the standard bonded/unbonded criterion used within the oxDNA framework. Melting probability profiles were obtained by calculating the fraction of sampled configurations in which each base pair was denatured.

To quantify the spatial organisation of melting events, we calculated the Gini coefficient of the melting probability distribution along each sequence. The Gini coefficient ranges from 0, indicating that melting probability is uniformly distributed across the sequence, to 1, indicating maximal inequality, with melting concentrated at a small number of positions. As a complementary measure we calculated the the Shannon entropy, *H* = − ∑*_i_ p_i_* log(*p_i_*),where *p_i_* denotes the probability that base pair *i* is melted. High Gini coefficients and low entropy values indicate that melting is concentrated at a small number of preferred sites, whereas lower Gini coefficients and higher entropy values reflect more spatially distributed melting.

#### Analysis of twist and writhe

The partitioning of imposed supercoiling into twist, writhe and denaturation was analysed from equilibrated DNA conformations. Local twist and writhe were calculated using standard geometric definitions of DNA topology. Representative molecular configurations were visualised using oxView and used to identify denaturation bubbles, plectonemic structures and regions of local writhe. In particular, simulations were analysed to determine the spatial relationship between localised melting events and plectoneme formation under negative supercoiling.

#### Simulations with nucleosome constraints

To investigate the influence of promoter chromatin architecture on supercoiling-induced melting, nucleosome occupancy information derived from Fiber-seq data was incorporated into the simulations. For each promoter, stochastic nucleosome configurations were generated by Monte Carlo sampling from experimentally measured nucleosome occupancy profiles.

Rather than explicitly modelling histone octamers, nucleosome-associated DNA segments were represented through effective mechanical constraints designed to reproduce the principal physical consequences of nucleosome occupancy. Specifically, nucleosomal DNA was assigned increased resistance to melting, bending and twisting deformations, corresponding to an increased effective persistence length and reduced torsional flexibility. These constraints strongly suppress local writhe formation and denaturation within nucleosome-bound regions while preserving the overall DNA connectivity and topology.

Independent nucleosome occupancy configurations were simulated using the same protocol described above. Melting profiles, twist and writhe distributions, and representative molecular conformations were subsequently compared between CGI and non-CGI promoters under equivalent levels of imposed supercoiling.

### Genomic datasets

Publicly available human genomic datasets used in this study were obtained from the NCBI Sequence Read Archive (SRA) and Gene Expression Omnibus (GEO) databases. Dataset accession numbers and corresponding references are provided in Table 1. Where processed genomic signal tracks were available, these were downloaded directly from the corresponding repository. For datasets obtained as raw sequencing reads, FASTQ files were downloaded from GEO/SRA and processed as described below. All genomic coordinates and datasets were analysed relative to the GRCh38/hg38 human reference genome; datasets originally mapped to alternative genome assemblies were converted to hg38 where required.

Published datasets were integrated with our Twist-seq data to examine the relationship between DNA supercoiling and genomic features including transcription, topoisomerase occupancy, DNA secondary structure and chromatin-associated features. Signal around transcription start sites was quantified using deepTools computeMatrix and visualised using plotProfile and with gene sets and genomic regions defined as described above.

### Transcription factor motif and DNA melting analysis

Transcription factor (TF) motifs were predicted within the 1-kb promoter sequences used for oxDNA simulations using FIMO (MEME Suite v5.5.9)36 and the JASPAR 2026 CORE vertebrate non-redundant motif collection37. Motif matches with a FIMO P value <1 × 10⁻⁴ were retained (Supplementary Table 2). FIMO coordinates were converted to the zero-based coordinate system used for the oxDNA simulations. Motif predictions were used as sequence annotations and were not interpreted as evidence of TF occupancy. To examine the local relationship between predicted TF motifs and DNA melting, the mean simulated melting probability within each predicted motif occurrence was compared with the immediately adjacent sequence. For each motif occurrence, the flanking value was calculated as the mean melting probability across the combined 25-bp regions immediately upstream and downstream of the motif. Δ melting probability was calculated as the mean melting probability within the motif minus the mean melting probability of these two 25-bp flanking regions. Thus, negative Δ values indicate that the motif-containing sequence has a lower mean melting probability than its immediately adjacent DNA. Motifs lacking sufficient flanking sequence at the ends of the simulated sequence were excluded. This analysis was performed across all 12 nucleosome-positioned promoter simulations. For each promoter, the Δ melting values from all eligible predicted motif occurrences were combined to generate a promoter-specific distribution, irrespective of TF identity. The median Δ melting probability was used to summarize the overall relationship between predicted TF motif-containing sequences and their local flanking DNA within each promoter.

The spatial relationship between predicted TF motifs and regions of strong DNA melting was examined further in the final nucleosome- and topologically constrained simulations of ATF4, EEF1A1, CCDC80 and RGS4. High-probability melting regions were defined as contiguous positions with a melting probability ≥0.1. This threshold was applied uniformly across promoter sequences and represents a high-probability melting threshold for the simulation profiles rather than the molecular criterion used within the simulations to classify an individual base pair as melted. For promoters containing high-probability melting regions, the proportion of predicted TF motif occurrences overlapping these regions was calculated. The observed overlap was compared with a null distribution generated from 10,000 within-promoter positional randomizations in which the observed number and length of motifs were preserved while their positions within the promoter were randomized. Empirical one-sided P values were calculated to test whether predicted TF motif overlap with high-probability melting regions was lower than expected under this positional null model.

### Somatic mutation analysis

PCAWG whole-genome somatic SNVs passing the ICGC consensus variant filters were obtained from the ICGC consensus mutation dataset^38^, lifted from GRCh37 to GRCh38 and restricted to primary chromosomes. The final dataset comprised 21,599,452 SNVs from 1,950 tumours across 25 PCAWG projects/cancer types. Mutation counts were quantified within ±500 bp of strand-aware transcription start sites for CGI and non-CGI promoters stratified by expression level. Somatic mutation counts were analysed using negative-binomial mixed-effects models, with promoter sequence opportunity and total tumour SNV burden included as offsets and tumour included as a random intercept. Rate ratios and 95% confidence intervals were estimated for CGI relative to non-CGI promoters within each expression group. An interaction between CGI status and expression group was tested using a likelihood-ratio test to determine whether the relative mutation rates of CGI and non-CGI promoters differed according to transcriptional activity.

## Data Availability

Raw and processed sequencing data generated in this study will be deposited in the Gene Expression Omnibus (GEO), and the accession number will be provided prior to publication.

## Acknowledgements

We thank all members of the Gilbert and Marenduzzo labs for useful discussions and critical reading of the manuscript, and Connor Warnock, IGC Graphics, for assistance with figure artwork. This work was funded by the UK Medical Research Council (MR/J00913X/1;MC_UU_00007/13)(NG) and Wellcome Trust investigator award 223097/Z/21 (MC, DMa, NG).

## Author contributions

C.N. J.S and S.C. undertook experiments, A.B, A.B., C.B., M.C. and D.M. carried out simulations. C.N., A.B., G.R.G., S.C., J.S., D.H., M.C. and D.M. analysed data. C.N., S.C., N.G. and D.M. conceived the project, and all authors contributed to writing the manuscript.

