## Supplemental Data for "CpG islands are topological sinks for transcription-induced DNA supercoiling"

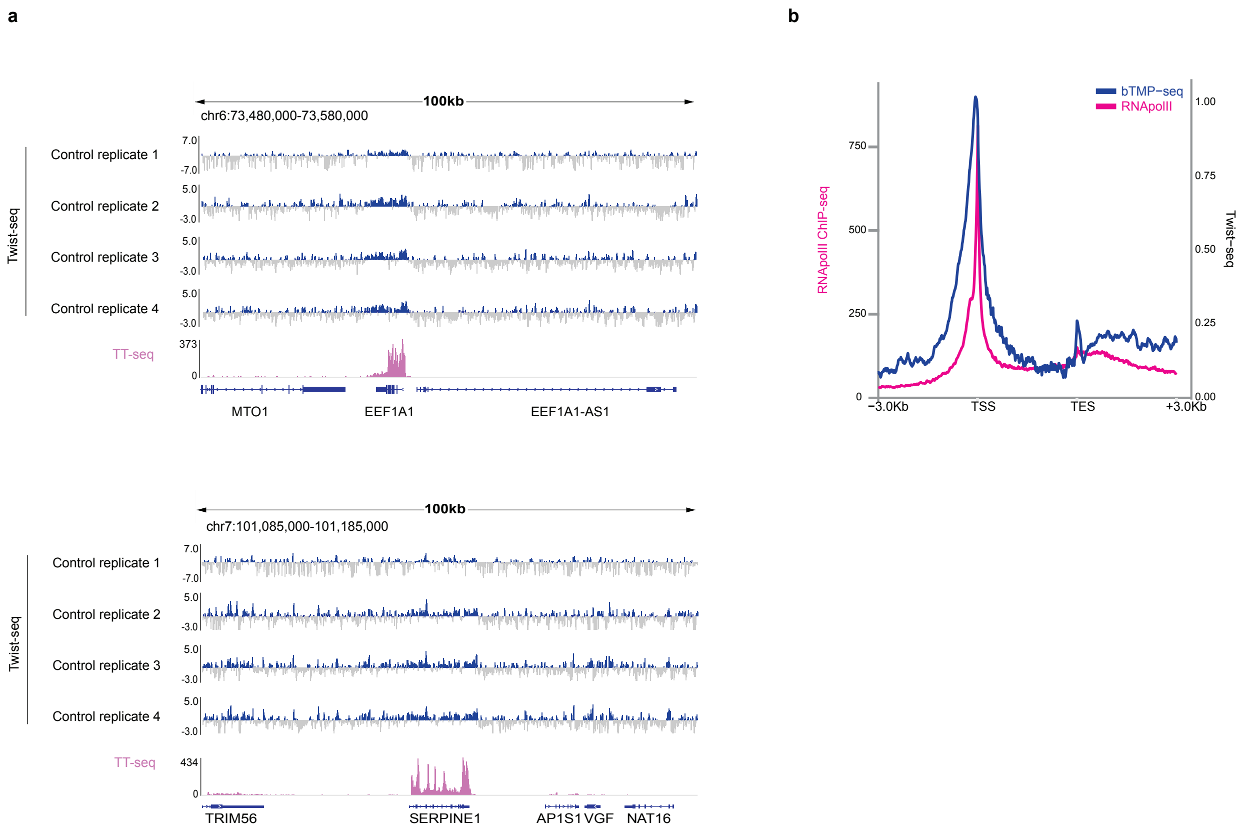
a
100kb
chr6:73,480,000-73,580,000
Twist-seq
Control replicate 1
Control replicate 2
Control replicate 3
Control replicate 4
TT-seq
7.0
-7.0
5.0
-3.0
373
0
MTO1
EEF1A1
EEF1A1-AS1
100kb
chr7:101,085,000-101,185,000
Twist-seq
Control replicate 1
Control replicate 2
Control replicate 3
Control replicate 4
TT-seq
434
TRIM56
SERPINE1
AP1S1
VGF
NAT16
b
bTMP−seq
RNApolII
RNApolII ChIP-seq
Twist−seq
750
500
250
0
1.00
0.75
0.50
0.25
0.00
−3.0Kb
TSS
TES
+3.0Kb

**Supplementary Figure 1: a,** Individual replicates of Twist-seq mapped genomic distribution of negative DNA supercoiling around transcriptionally the active genes EEF1A1 and SERPINE. Twist-seq data is log2(Twist-seq control/Twist-seq genomic) in 1 bp bins. TT-seq mapped nascent gene expression is shown as RPKM. **b,** DNA supercoiling levels (Twist-seq) and RNA polymerase II Chip-seq[18] averaged across all genes within the high expression group with a 3-kb extension on both sides.

a

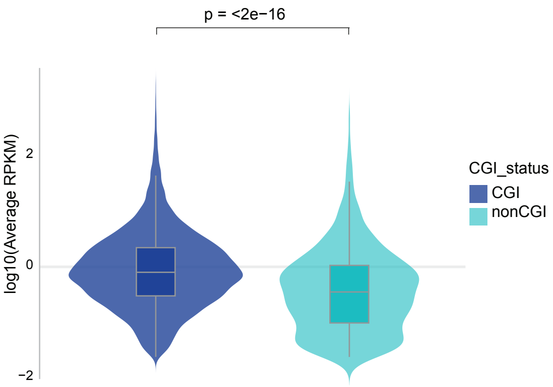
p = <2e−16
log10(Average RPKM)
2
0
−2
CGI_status
CGI
nonCGI

b

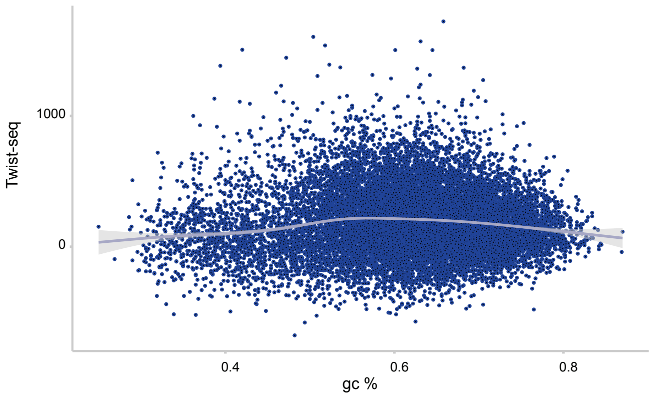
Twist-seq
1000
0
0.4
0.6
0.8
gc %

c

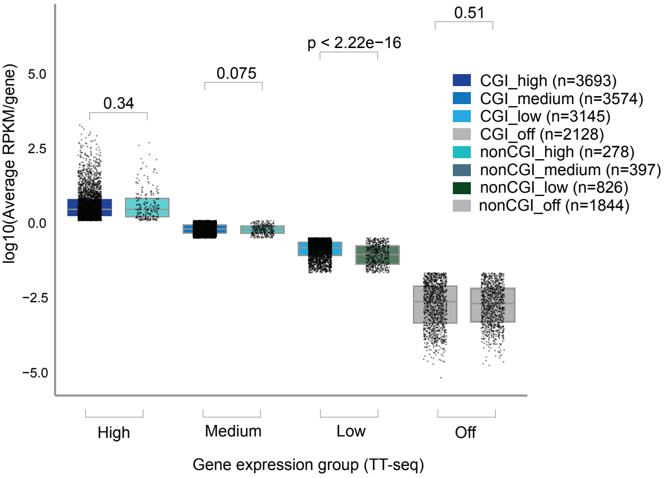
log10(Average RPKM/gene)
5.0
2.5
0.0
−2.5
−5.0
0.34
0.075
p < 2.22e−16
0.51
CGI_high (n=3693)
CGI_medium (n=3574)
CGI_low (n=3145)
CGI_off (n=2128)
nonCGI_high (n=278)
nonCGI_medium (n=397)
nonCGI_low (n=826)
nonCGI_off (n=1844)
High
Medium
Low
Off
Gene expression group (TT-seq)

d

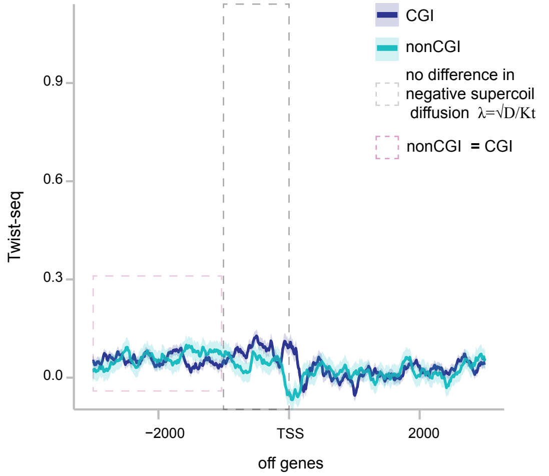
CGI
nonCGI
no difference in
negative supercoil
diffusion λ=√D/Kt
nonCGI = CGI
Twist-seq
0.9
0.6
0.3
0.0
−2000
TSS
2000
off genes

e

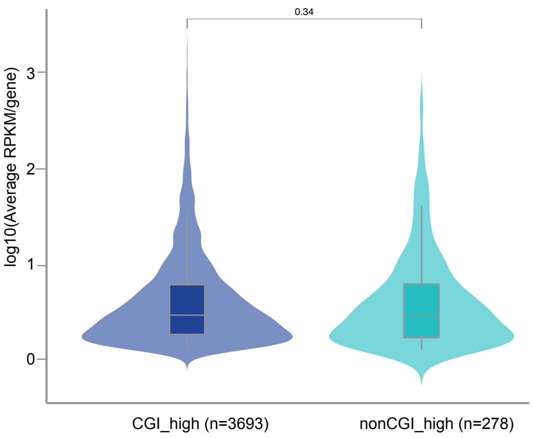
0.34
log10(Average RPKM/gene)
3
2
1
0
CGI_high (n=3693)
nonCGI_high (n=278)

f

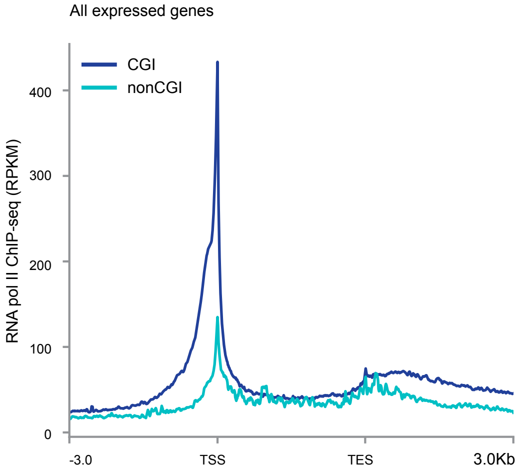

All expressed genes
CGI
nonCGI
RNA pol II ChIP-seq (RPKM)
400
300
200
100
0
-3.0
TSS
TES
3.0Kb


g

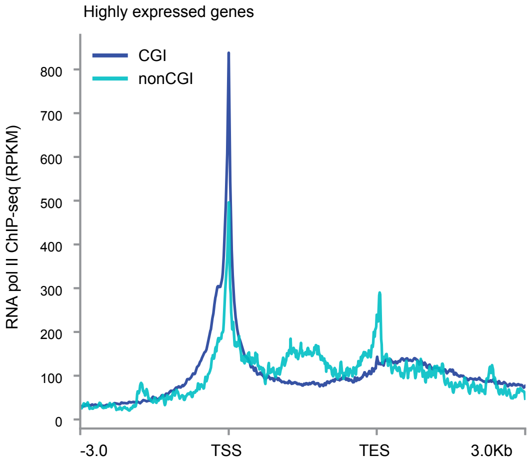

Highly expressed genes
CGI
nonCGI
RNA pol II ChIP-seq (RPKM)
800
700
600
500
400
300
200
100
0
-3.0
TSS
TES
3.0Kb

**Supplementary Figure 2: a,** Distribution of gene expression levels as measured by TT-seq (log10(average RPKM/gene)) at CGI and non-CGI promoter genes. *P* value was calculated using nonparametric Wilcoxon two-sided test. **b,** Correlation between the distribution of DNA supercoiling (Twist-seq +/-500 bp at TSS) and DNA sequence GC% (+/-500 bp at TSS) at gene promoters. **c**, Gene expression (TT-seq average RPKM/gene) distribution split by expression group (off, low, medium and high) and CGI status (CGI or non-CGI). *P* values were calculated using nonparametric Wilcoxon two-sided tests. **d**, DNA supercoiling levels (Twist-seq, 1bp bins) ±3 kb around TSSs, averaged across all genes within the off expression group for CGI and non-CGI promoters. **e,** Distribution of gene expression levels, for genes in the high expression group, as measured by TT-seq (log10(average RPKM/gene)) at CGI and non-CGI promoter genes. *P* value was calculated using nonparametric Wilcoxon two-sided test. **f,** RNA polymerase II (RPKM) levels averaged across all genes split by promoter CGI status. **g,** RNA polymerase II (RPKM) levels averaged across all genes within the high expression group and split by promoter CGI status.

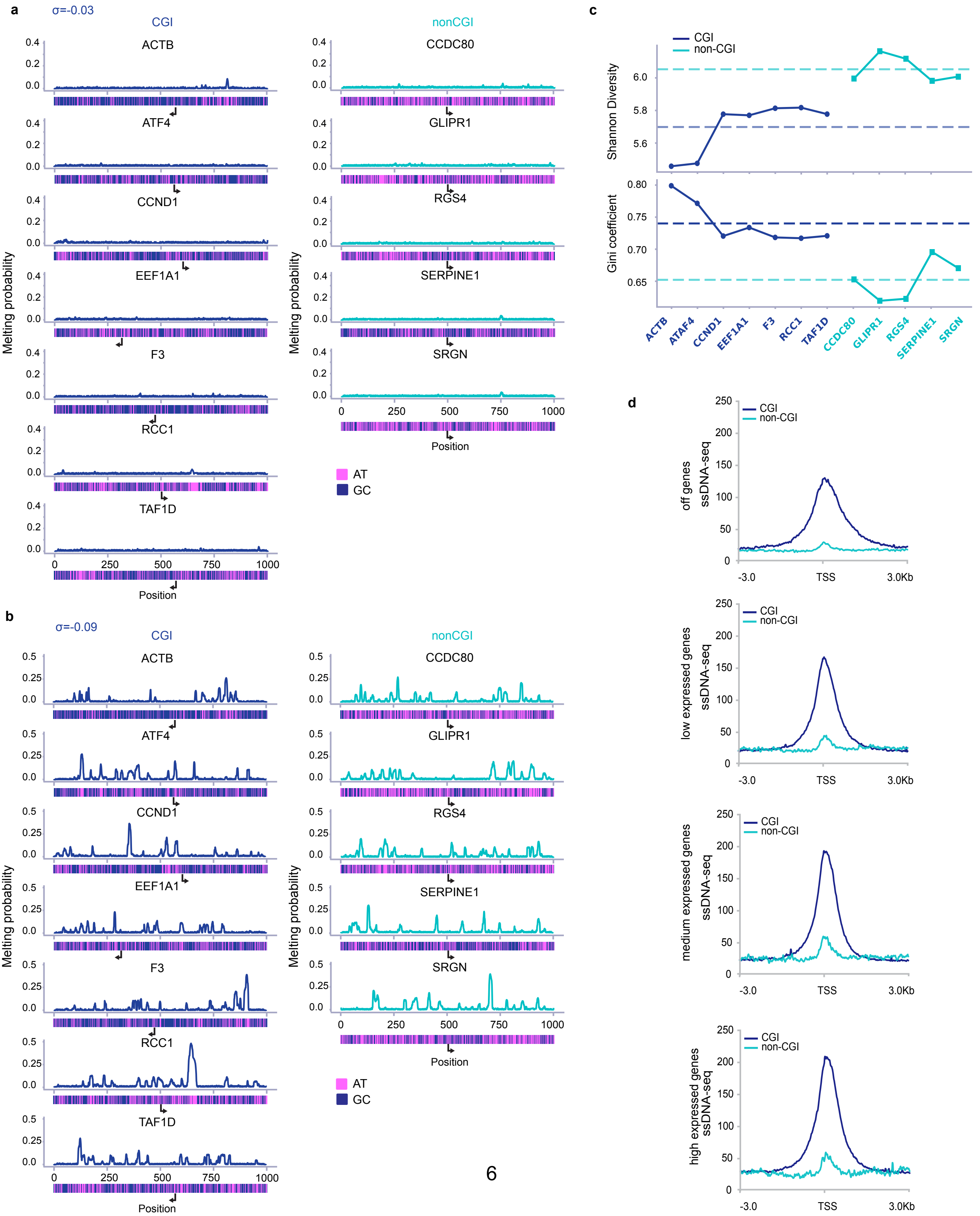

a
σ=-0.03
CGI
nonCGI
ACTB
ATF4
CCND1
EEF1A1
F3
RCC1
TAF1D
CCDC80
GLIPR1
RGS4
SERPINE1
SRGN
Melting probability
Position
AT
GC
b
σ=-0.09
c
CGI
non-CGI
Shannon Diversity
Gini coefficient
d
off genes ssDNA-seq
low expressed genes ssDNA-seq
medium expressed genes ssDNA-seq
high expressed genes ssDNA-seq
-3.0
TSS
3.0Kb

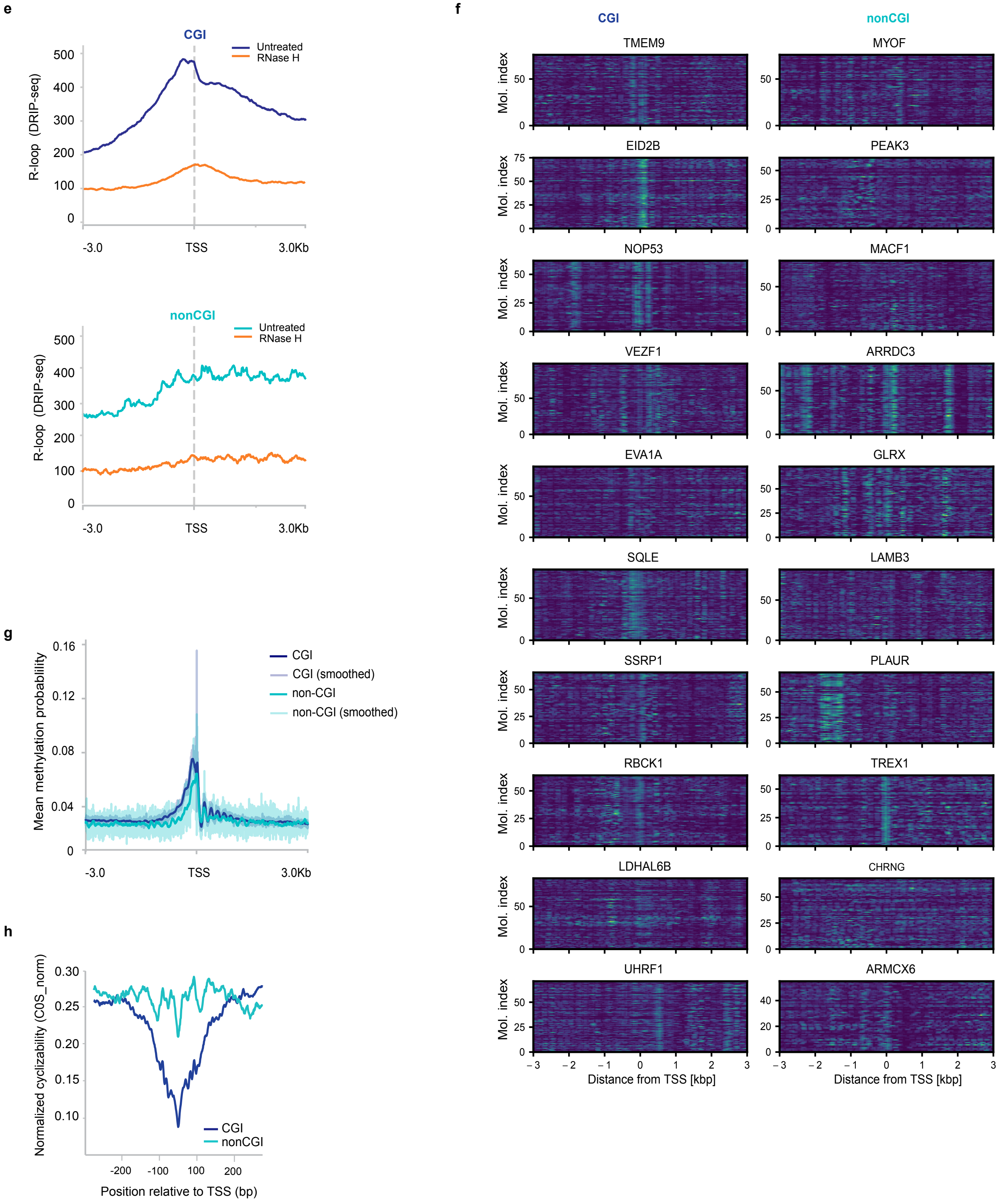
e
CGI
Untreated
RNase H
R-loop (DRIP-seq)
500
400
300
200
100
0
-3.0
TSS
3.0Kb
nonCGI
Untreated
RNase H
R-loop (DRIP-seq)
-3.0
TSS
3.0Kb
f
CGI
nonCGI
TMEM9
MYOF
EID2B
PEAK3
NOP53
MACF1
VEZF1
ARRDC3
EVA1A
GLRX
SQLE
LAMB3
SSRP1
PLAUR
RBCK1
TREX1
LDHAL6B
CHRNG
UHRF1
ARMCX6
Mol. index
Distance from TSS [kbp]
Distance from TSS [kbp]
g
Mean methylation probability
0.16
0.12
0.08
0.04
0
CGI
CGI (smoothed)
non-CGI
non-CGI (smoothed)
-3.0
TSS
3.0Kb
h
Normalized cyclizability (C0S_norm)
0.30
0.25
0.20
0.15
0.10
CGI
nonCGI
-200
-100
100
200
Position relative to TSS (bp)

**Supplementary Figure 3: a, b,** Melting profile for CGI (left panel) and non-CGI (right panel) promoter DNA sequences with a supercoil density of σ=-0.03 (a) and σ=-0.09 (b). GC/AT content heatmaps are shown under each simulation and arrows highlight the position and direction of TSS. **c**, Quantification of the spatial distribution of DNA melting in CGI and non-CGI promoter sequences at σ = −0.06 using the Gini coefficient and Shannon entropy. Higher Gini coefficients and lower Shannon entropy indicate more spatially localised melting. **d,** ssDNA-seq[18] signal (RPKM, 1 bp bins) averaged across ±3 kb around TSSs of CGI and non-CGI genes split by expression group (off, low, medium and high expressed genes)**. e,** Mean DRIP-seq signal from published RPE1 data is shown across highly expressed CGI (upper panel) and non-CGI promoters (lower panel) centred on the TSS. **f,** Single molecule nucleosome density plots of Fiber-seq data +/-3Kb TSS for CGI and nonCGI genes (ref). **g**, Fiber-seq mean methylation probability (50 bp bins) averaged across ±3 kb around TSSs of CGI and non-CGI genes in expression group high. **h**, DNAcycP prediction of DNA cyclizability for DNA sequences in a 2kb window at TSS for CGI versus non-CGI promoters[56]**.**

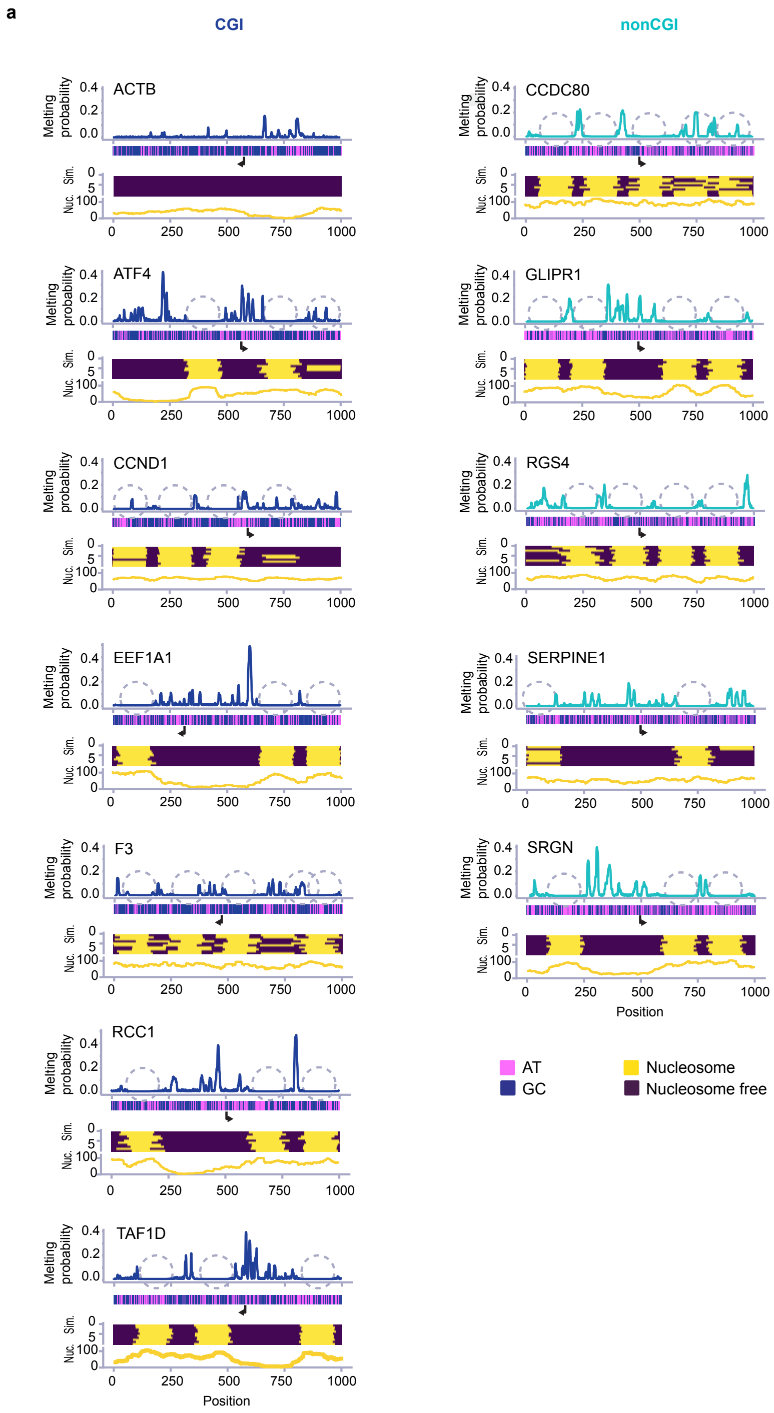
a
CGI
nonCGI
ACTB
ATF4
CCND1
EEF1A1
F3
RCC1
TAF1D
CCDC80
GLIPR1
RGS4
SERPINE1
SRGN
Melting probability
Sim.
Nuc.
Position
AT
GC
Nucleosome
Nucleosome free

b

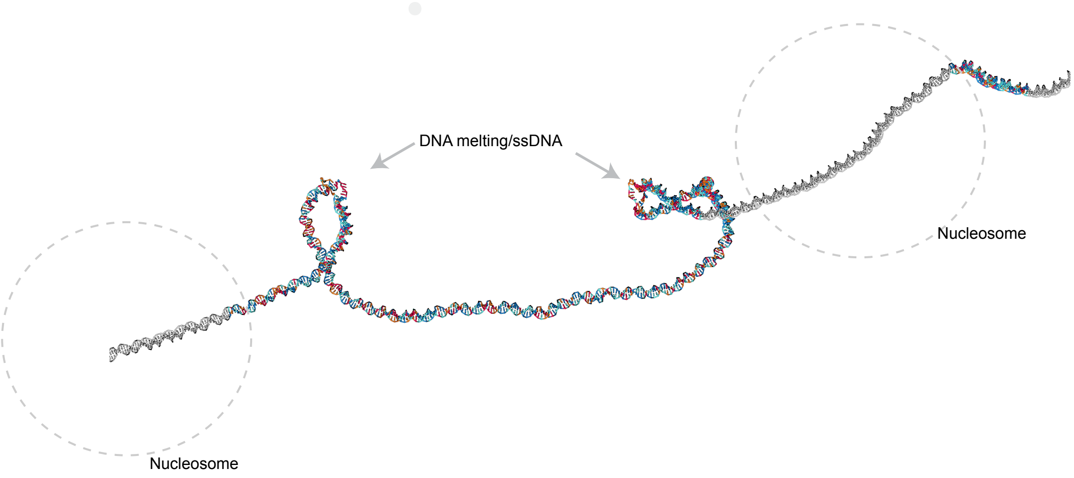

DNA melting/ssDNA
Nucleosome
Nucleosome


c

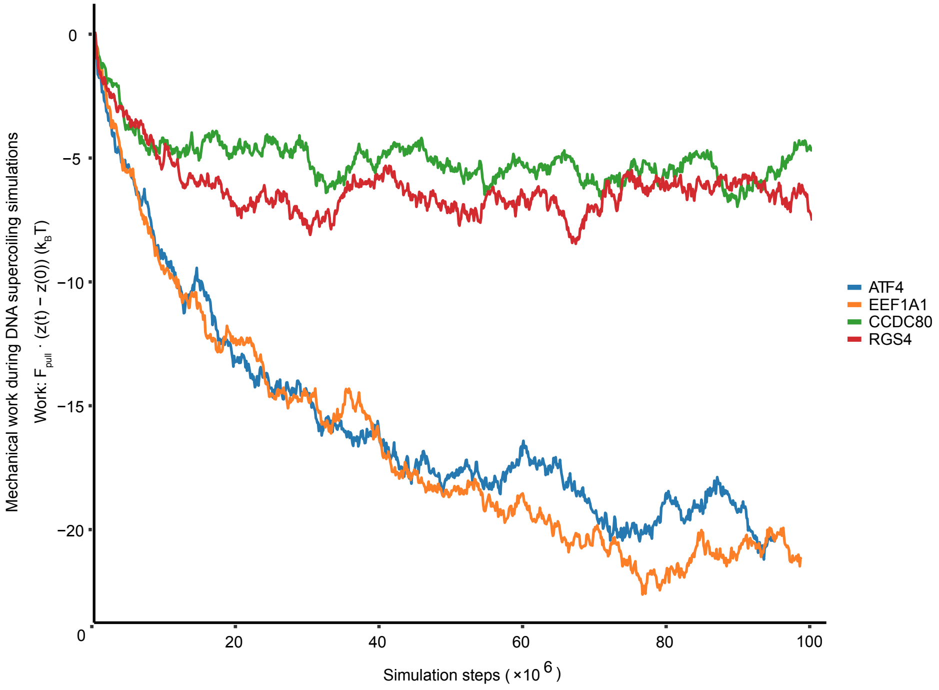

Mechanical work during DNA supercoiling simulations
Work: $F_{pull} \cdot (z(t) - z(0))$ ($k_BT$)
0
−5
−10
−15
−20
0
20
40
60
80
100
Simulation steps ( ×10 6 )
ATF4
EEF1A1
CCDC80
RGS4

**Supplementary Figure 4: a,** Melting profile for CGI and non-CGI promoter DNA sequences with supercoil density of σ=-0.06 and nucleosomes positioned as indicated by the dashed circles. Nucleosome positioning was predicted from Monte Carlo based simulations (Sim track) of fibre seq data (Nuc track). **b,** Simulation snapshot illustrating DNA writhe and melting at EEF1A1 CGI promoter. Mechanical work calculated during simulations of negatively supercoiled DNA (σ = −0.06) for two CGI (ATAF4 and EEF1A1) and two non-CGI (CCDC80 and RGS4) promoter sequences. **c,** Mechanical work calculated during simulations of negatively supercoiled DNA (σ = −0.06) for two CGI (ATAF4 and EEF1A1) and two non-CGI (CCDC80 and RGS4) promoter sequences. Work was calculated as the applied pulling force (Fpull) multiplied by the change in DNA extension relative to its initial value (z(t) − z(0)). Work is expressed in units of kBT and plotted against simulation step.

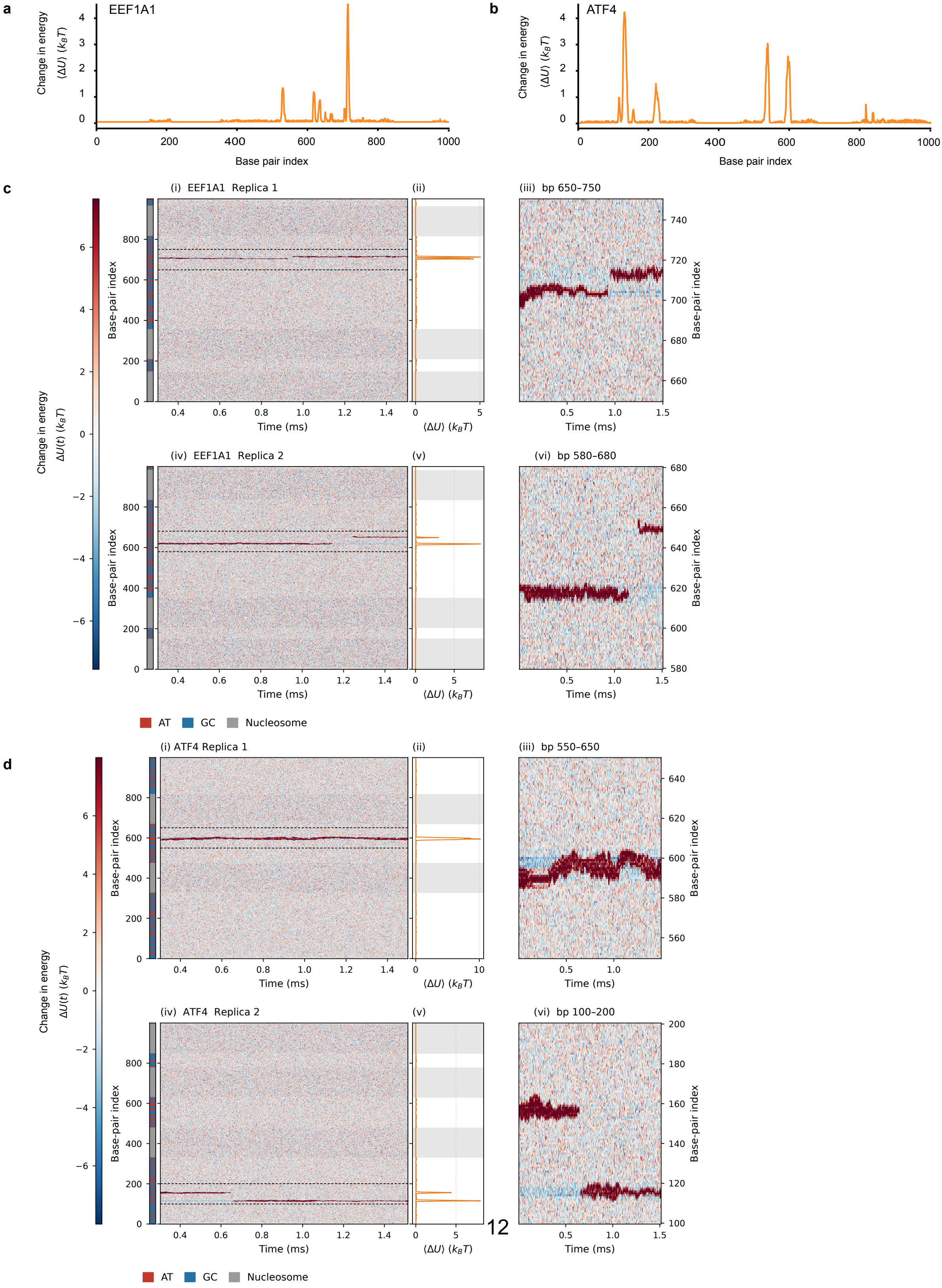

a
EEF1A1
Change in energy
⟨ΔU⟩ ($k_BT$)
Base pair index
b
ATF4
c
(i) EEF1A1 Replica 1
(ii)
(iii) bp 650–750
Change in energy
ΔU(t) ($k_BT$)
Base-pair index
Time (ms)
⟨ΔU⟩ ($k_BT$)
(iv) EEF1A1 Replica 2
(v)
(vi) bp 580–680
AT
GC
Nucleosome
d
(i) ATF4 Replica 1
(iii) bp 550–650
(iv) ATF4 Replica 2
(vi) bp 100–200

**Supplementary Figure 5: Sequence-dependent DNA melting and associated internal energy changes in oxDNA simulations incorporating nucleosome occupancy and mechanical constraints. a,b,** Mean internal energy difference ($\langle \Delta U \rangle$) along the EEF1A1 (**a**) and ATF4 (**b**) CGI promoter sequences in simulations incorporating sequence-dependent nucleosome occupancy and mechanical constraints on nucleosome-bound DNA. The internal energy difference was calculated as the difference between the mean internal energy and the mean energy of the bonded (hybridised) states, averaged across all simulation replicas. Values are expressed in units of kBT. **c,d,** Time-resolved DNA melting and associated internal energy changes for EEF1A1 (**c**) and ATF4 (**d**). For each promoter, two representative simulation replicas are shown. Panels (i) and (iv) show the internal energy difference ($\Delta U$) over time for two individual simulation replicas. Adjacent panels (ii) and (v) show the corresponding time-averaged internal energy differences along the sequence for the same individual replicas. Panels (iii) and (vi) show magnified views of the regions indicated by dashed boxes in the kymographs, highlighting the spatial and temporal distribution of DNA melting. Sequence annotations indicate AT and GC base pairs and nucleosome-associated regions. Time is expressed in milliseconds using the simulation-to-physical-time mapping described in Methods.

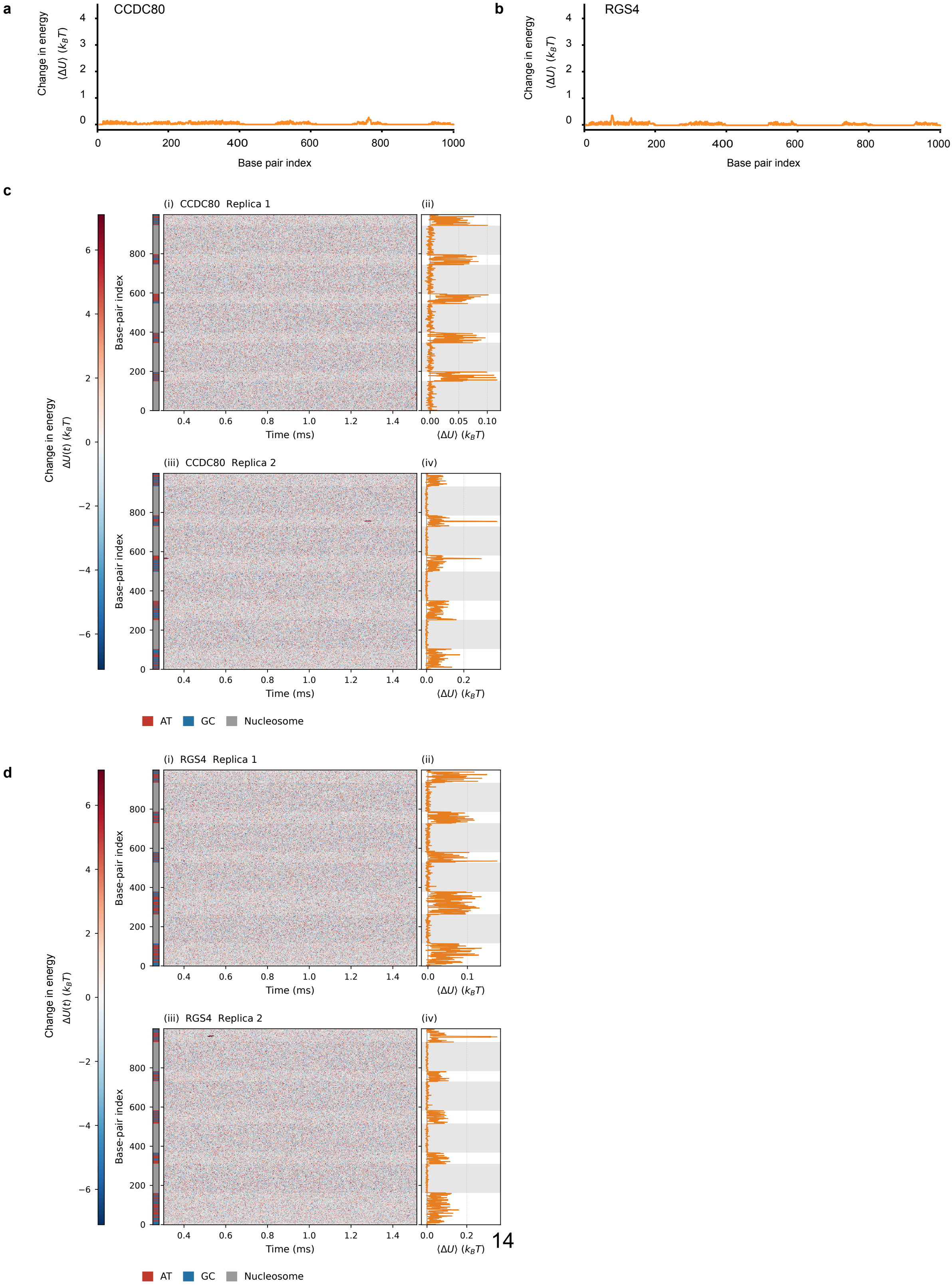

a
CCDC80
Change in energy
⟨ΔU⟩ ($k_BT$)
Base pair index
b
RGS4
Change in energy
⟨ΔU⟩ ($k_BT$)
Base pair index
c
(i) CCDC80 Replica 1
(ii)
(iii) CCDC80 Replica 2
(iv)
Change in energy
ΔU(t) ($k_BT$)
Base-pair index
Time (ms)
⟨ΔU⟩ ($k_BT$)
AT
GC
Nucleosome
d
(i) RGS4 Replica 1
(ii)
(iii) RGS4 Replica 2
(iv)
Change in energy
ΔU(t) ($k_BT$)
Base-pair index
Time (ms)
⟨ΔU⟩ ($k_BT$)
AT
GC
Nucleosome

**Supplementary Figure 6: Internal energy changes in oxDNA simulations of non-CGI promoter sequences incorporating nucleosome occupancy and mechanical constraints. a,b,** Mean internal energy difference (⟨ΔU⟩) along the CCDC80 (**a**) and RGS4 (**b**) non-CGI promoter sequences in simulations incorporating sequence-dependent nucleosome occupancy and mechanical constraints on nucleosome-bound DNA. The internal energy difference was calculated as the difference between the mean internal energy and the mean energy of the bonded (hybridised) states, averaged across all simulation replicas. Values are expressed in units of kBT. **c,d,** Time-resolved internal energy changes for CCDC80 (**c**) and RGS4 (**d**). For each promoter, two representative simulation replicas are shown. Panels (i) and (iii) show kymographs of the internal energy difference (ΔU) over time for two individual simulation replicas. Adjacent panels (ii) and (iv) show the corresponding time-averaged internal energy differences along the sequence for the same individual replicas. Sequence annotations indicate AT and GC base pairs and nucleosome-associated regions. Time is expressed in milliseconds using the simulation-to-physical-time mapping described in Methods. Note: The horizontal scales of the time-averaged internal energy profiles in c and d differ from those in the corresponding CGI promoter simulations (Supplementary Fig. 5), reflecting the substantially smaller internal energy differences observed for the non-CGI sequences.

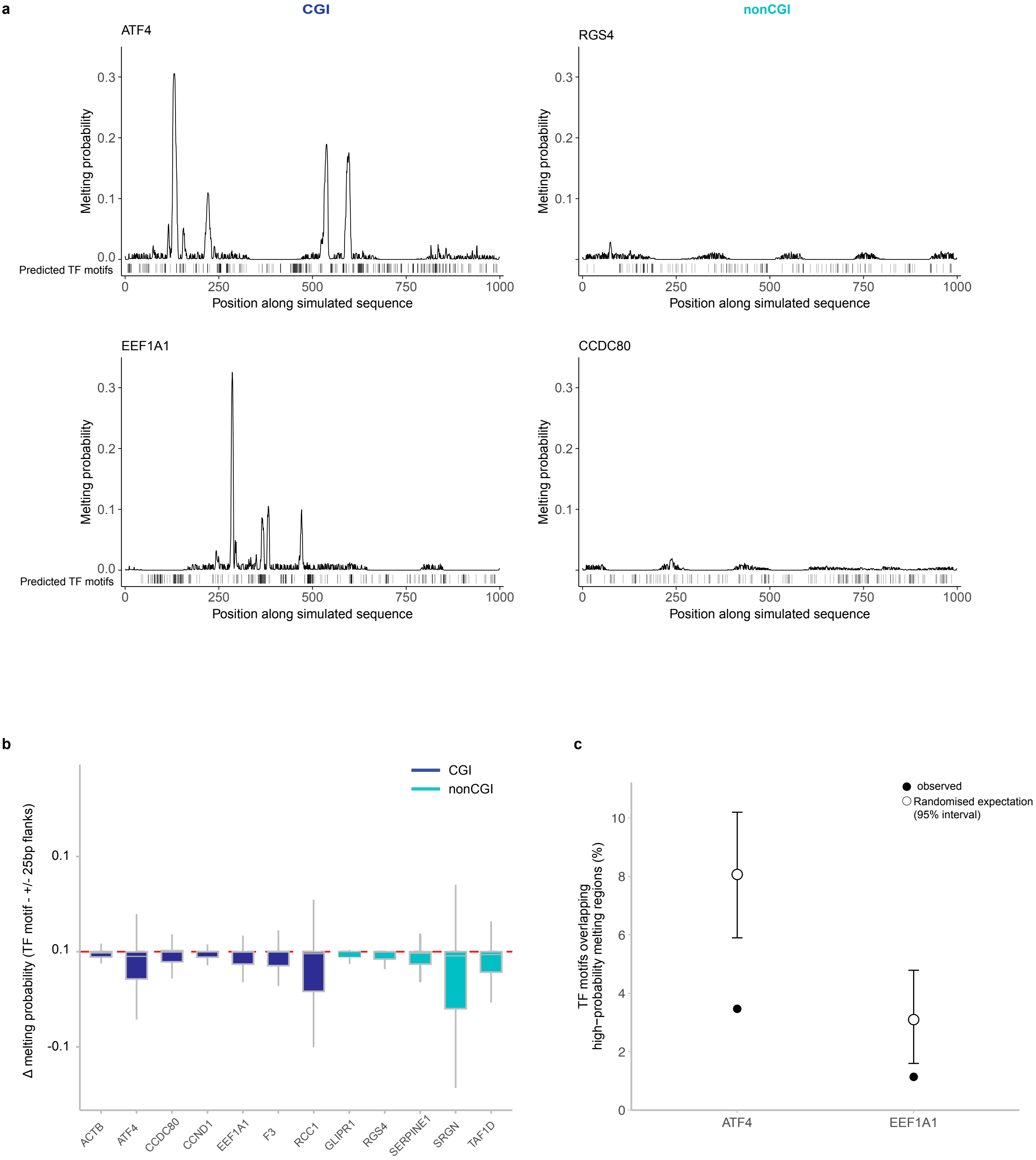
a
CGI
nonCGI
ATF4
RGS4
EEF1A1
CCDC80
Melting probability
Predicted TF motifs
Position along simulated sequence
0.0
0.1
0.2
0.3
0
250
500
750
1000
b
CGI
nonCGI
Δ melting probability (TF motif - +/- 25bp flanks)
0.1
0.1
-0.1
ACTB
ATF4
CCDC80
CCND1
EEF1A1
F3
RCC1
GLIPR1
RGS4
SERPINE1
SRGN
TAF1D
c
observed
Randomised expectation
(95% interval)
TF motifs overlapping
high−probability melting regions (%)
0
2
4
6
8
10
ATF4
EEF1A1

**Supplementary Figure 7: a,** DNA melting profiles and predicted TF motif positions in nucleosome- and topologically constrained promoter simulations. Simulated melting probability is shown across the 1-kb promoter sequences for ATF4, RGS4, EEF1A1 and CCDC80. Vertical ticks beneath each profile indicate the positions of predicted TF motif occurrences identified by FIMO, irrespective of TF identity. **b,** Local DNA melting probability at predicted TF motifs across 12 nucleosome-positioned promoter simulations. For each predicted TF motif occurrence, Δ melting probability was calculated as the mean melting probability within the motif minus the mean melting probability across the combined 25-bp regions immediately upstream and downstream. Negative values therefore indicate lower melting probability within the motif-containing sequence than in its immediately adjacent DNA. Each boxplot shows the distribution of Δ melting values for all eligible predicted motif occurrences within that promoter, irrespective of TF identity. **c,** Predicted TF motif overlap with high-probability melting regions in the constrained promoter simulations. Filled circles indicate the observed proportion of predicted TF motifs overlapping high-probability melting regions (melting probability ≥0.1); open circles and error bars indicate the mean and 95% interval obtained from 10,000 within-promoter positional randomizations. Predicted TF motifs were significantly depleted from high-probability melting regions in both ATF4 ($P = 1 \times 10^{-4}$) and EEF1A1 ($P = 0.0068$).

**a**

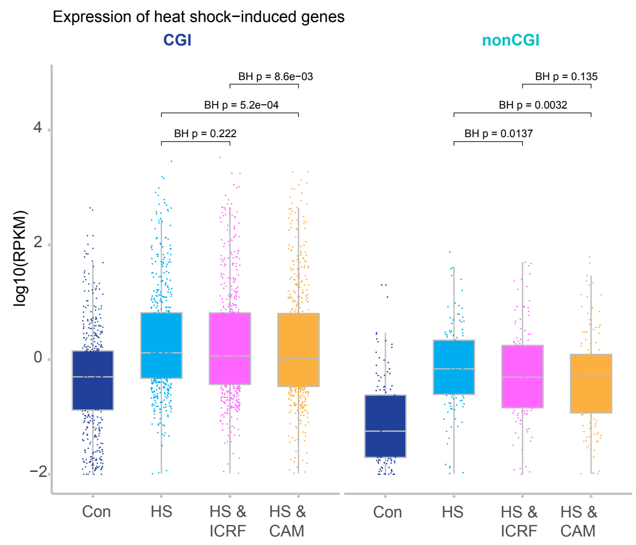

Expression of heat shock−induced genes
CGI
nonCGI
BH p = 8.6e−03
BH p = 5.2e−04
BH p = 0.222
BH p = 0.135
BH p = 0.0032
BH p = 0.0137
log10(RPKM)
4
2
0
−2
Con
HS
HS & ICRF
HS & CAM
Con
HS
HS & ICRF
HS & CAM


**b**

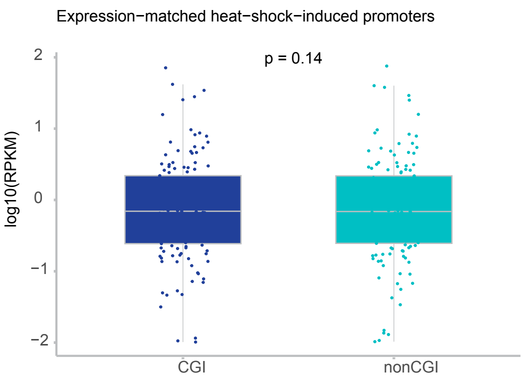

Expression−matched heat−shock−induced promoters
p = 0.14
log10(RPKM)
2
1
0
−1
−2
CGI
nonCGI

**Supplementary Figure 8: a,** Expression of heat-shock-induced CGI (n = 523) and non-CGI (n = 132) genes measured by TT-seq under control (CON), heat shock (HS), TOP2 inhibition during heat shock (HS & ICRF) and TOP1 inhibition during heat shock (HS & CAM) conditions. Boxplots show gene expression ($\log_{10}$(RPKM + 0.01)). Individual points represent genes; boxes indicate the median and interquartile range. Pairwise comparisons were performed using paired two-sided Wilcoxon signed-rank tests with Benjamini–Hochberg correction. For CGI genes, HS versus HS & ICRF, adjusted P = 0.222; HS versus HS & CAM, adjusted P = $5.2 \times 10^{-4}$; and HS & ICRF versus HS & CAM, adjusted P = $8.6 \times 10^{-3}$. For non-CGI genes, HS versus HS & ICRF, adjusted P = 0.0137; HS versus HS & CAM, adjusted P = 0.0032; and HS & ICRF versus HS & CAM, adjusted P = 0.135. **b,** Expression matching of heat-shock-induced CGI and non-CGI genes. Boxplots show gene expression ($\log_{10}$(RPKM + 0.01)) following heat shock for expression-matched CGI and non-CGI gene sets (n = 129 per group; Supplementary Table 4). Individual points represent genes; boxes indicate the median and interquartile range. Expression did not differ significantly between the matched groups (paired two-sided Wilcoxon signed-rank test, P = 0.14).

**Supplementary Table 1:** Published genomic datasets used in this study, including experiment type, accession number, human cell line and corresponding reference.

**Supplementary Table 2:** Predicted transcription factor motifs in promoter sequences used for DNA melting simulations. TF motif sites were identified using FIMO (MEME Suite)[36] with the JASPAR 2026 CORE vertebrate non-redundant motif collection[37] and a motif-match threshold of $P < 1 \times 10^{-4}$. The Summary sheet reports the total number of predicted TF motif sites in each promoter, the number included in the analysis comparing melting probability within motif-containing sequences with the immediately flanking 25-bp sequences, and the number of distinct TF motifs represented. The Predicted TF motif sites sheet lists all 4,940 individual motif predictions, including motif identity, JASPAR ID, position, strand, matched sequence, FIMO score, P value and q value. Motif coordinates are reported relative to the simulated promoter sequence, with position 0 corresponding to the first nucleotide.

**Supplementary Table 3: Heat-shock-induced genes identified by TT-seq.** Genes exhibiting increased transcription following heat shock, identified using an independent TT-seq dataset. Heat-shock-induced genes were defined by a $\log_2$ fold change > 1 and an adjusted P value < 0.05. The table provides Ensembl gene identifiers, gene symbols, CGI promoter classification and differential expression statistics, including mean normalised counts, $\log_2$ fold changes, standard errors, test statistics and P values. This gene set was used to identify heat-shock-induced promoters for the Twist-seq analyses shown in Fig. 7b,c.

**Supplementary Table 4**: List of 129 CGI and 129 non-CGI genes matched according to their expression levels following heat shock, as measured by TT-seq. Non-CGI genes were ordered by increasing log-transformed expression ($\log_{10}$(RPKM + 0.01)) and paired with the CGI gene exhibiting the closest expression level, without replacement. The table provides Ensembl gene identifiers, gene symbols and expression values (RPKM and $\log_{10}$(RPKM + 0.01)) for each matched pair. These gene sets were used for the Twist-seq analyses shown in Fig. 7d,e and the expression comparison in Supplementary Fig. 8b.
